# Mapping lineage-traced cells across time points with moslin

**DOI:** 10.1101/2023.04.14.536867

**Authors:** Marius Lange, Zoe Piran, Michal Klein, Bastiaan Spanjaard, Dominik Klein, Jan Philipp Junker, Fabian J. Theis, Mor Nitzan

## Abstract

Simultaneous profiling of single-cell gene expression and lineage history holds enormous potential for studying cellular decision-making beyond simpler pseudotime-based approaches. However, it is currently unclear how lineage and gene expression information across experimental time points can be combined in destructive experiments, which is particularly challenging for in-vivo systems. Here we present moslin, a Fused Gromov-Wasserstein-based model to couple matching cellular profiles across time points. In contrast to existing methods, moslin leverages both intra-individual lineage relations and inter-individual gene expression similarity. We demonstrate on simulated and real data that moslin outperforms state-of-the-art approaches that use either one or both data modalities, even when the lineage information is noisy. On *C. elegans* embryonic development, we show how moslin, combined with trajectory inference methods, predicts fate probabilities and putative decision driver genes. Finally, we use moslin to delineate lineage relationships among transiently activated fibroblast states during zebrafish heart regeneration. We anticipate moslin to play a crucial role in deciphering complex state change trajectories from lineage-traced single-cell data.

## Introduction

Important biological processes like development, disease, or regeneration play out as complex changes on the cellular level. Due to their transforming nature, these changes are best captured by time-resolved measurements. Single-cell assays, including single-cell RNA-sequencing (scRNA-seq), probe cellular heterogeneity at unprecedented resolution and scale at different time points but destroy cells in the process. Thus, previous work introduced computational approaches that link cells across time based on similar gene expression profiles^1–3^. While these approaches successfully uncovered trajectories and fate decisions for in-vitro systems^1,4^ and some in-vivo systems^5,6^, they require dense temporal sampling and remain limited to simpler processes where expression similarity faithfully represents lineage relationships^7^.

To improve the accuracy of trajectory inference, scRNA-seq has been combined with heritable barcodes that link clonally related cells over long time scales in single-cell lineage tracing (scLT) assays^8–13^. For in-vitro systems, we can sample from the same cell population several times, and previous methods used this setting to relate cells across time points clonally^14,15^. However, such strategies do not generalize to in-vivo lineage-traced systems, as each time point corresponds to a different individual, and barcodes are not comparable across individuals. Most current analysis strategies^13,16–20^ remain limited to analyzing isolated lineage-traced time points. Thus, they do not embed lineage relationships in the temporal context of cellular state changes.

While a previous method, LineageOT^21^, represented an important step towards mapping lineage-traced cells, it cannot relate lineage information across time points and includes it only in the later time point. Further, the tool has only been demonstrated on simulated examples or examples with known ground truth. Thus, the comprehensive integration of lineage and gene expression information to estimate cellular state-change trajectories remains an open computational problem.

Here, we present multi-omic single-cell optimal transport for lineage data (moslin), a computational method to embed in-vivo clonal dynamics in their temporal context. Moslin uses expression similarity and lineage concordance to reconstruct cellular state-change trajectories for complex biological processes. To the best of our knowledge, moslin is the first method to use lineage information at two or more time points and to include the effects of cellular growth and stochastic cell sampling. Our approach outperforms LineageOT and optimal transport (OT)-baselines on simulated data where ground truth is available. Further, on *Caenorhabditis (C*.*) elegans* embryogenesis, we combine moslin with CellRank^22^, a trajectory inference framework, to uncover differentiation trajectories and putative decision-driver genes. Finally, in zebrafish heart regeneration, we predict lineage relationships between recently discovered activated fibroblast states that emerge after injury using moslin. We implemented moslin as a user-friendly Python package with documentation and tutorials, available at github.com/theislab/moslin.

## Results

### Moslin combines lineage and state information to link cells across time

Moslin is an algorithm to reconstruct molecular trajectories of complex cellular state changes from time-series single-cell lineage tracing^8,23,24^ (scLT) studies. Using gene expression and lineage information, moslin computes probabilistic mappings between cells in early- and late time points. Moslin distinguishes itself from previous approaches^21^ by incorporating lineage information at both time points to guide the inference process. Using the computed mapping, we infer ancestor and descendant probabilities for rare or transient cell states and interface with CellRank^22^ to visualize gene expression trends, uncover activation cascades, and pinpoint potential regulators of key decision events (Fig. 1a).

**Fig. 1.**
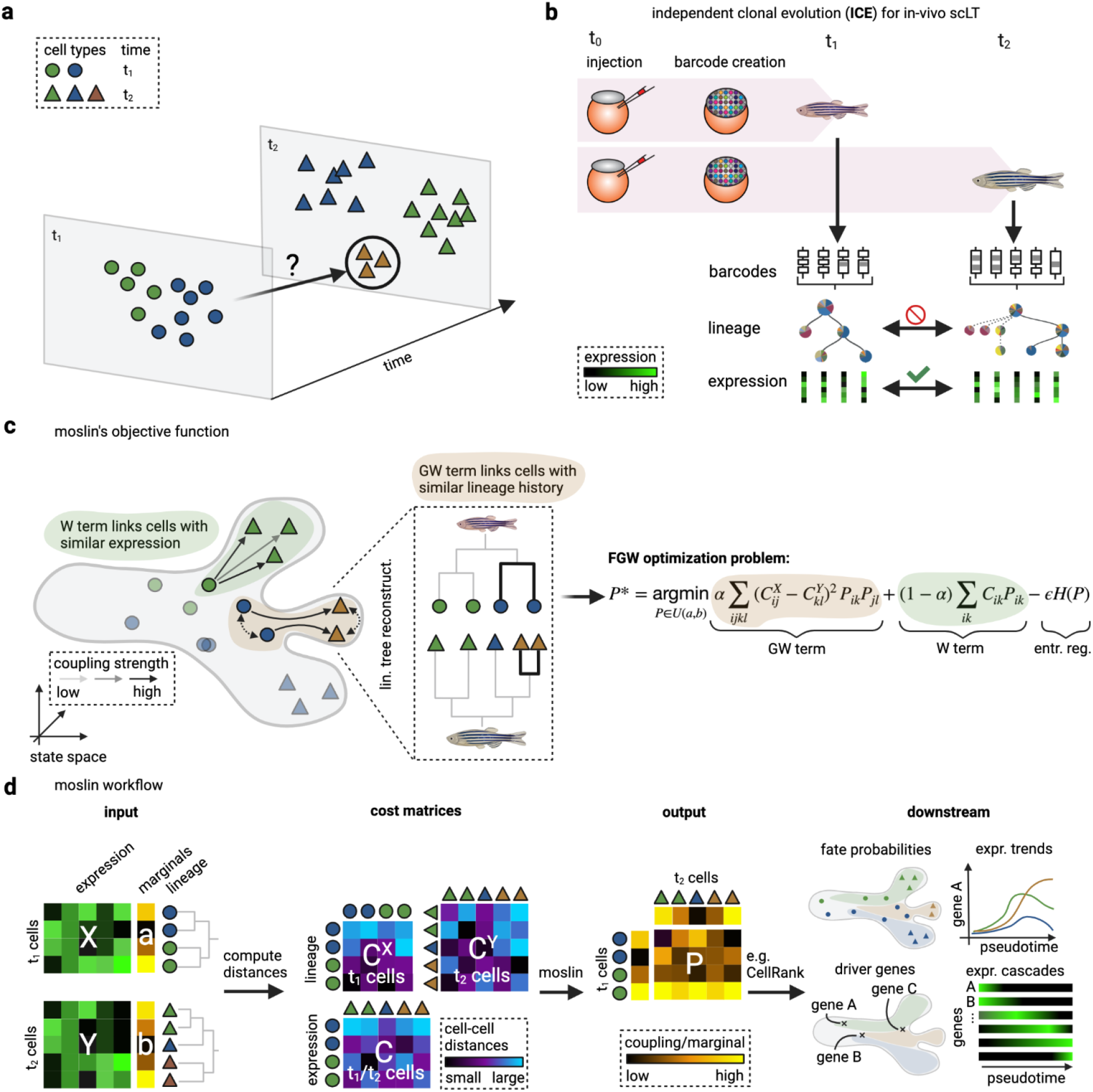
Moslin maps lineage-traced single cells across time points. **a**. Schematic of scRNA-seq time-course experiment with time points t_1_ (circles) and t_2_ (triangles). Cells are destroyed upon sequencing; this makes it difficult to study the trajectories of early cells giving rise to late cells. We highlight a rare population (brown triangles) that only appears at t_2_, with uncertain origin at t_1_. **b**. Illustration of independent clonal evolution (ICE) experimental design for scLT studies. ICE samples cells from different individuals at different time points and is applicable to in-vivo settings. **c**. Overview of moslin’s optimal-transport (OT)-based objective function for in-vivo scLT. The grey outline shows a simplified state manifold; shapes and colors as in (**a**). The dashed inset highlights lineage trees reconstructed independently for each time point^16^; these trees may be used in moslin to quantify lineage similarity. We use Wasserstein (W) and Gromov-Wasserstein (GW)-terms to compare cells in terms of gene expression and lineage similarity, respectively. The combination of W- and GW-terms gives rise to moslin’s Fused Gromow-Wasserstein (FGW) objective function on the right (Methods). **d**. The moslin workflow; based on gene expression matrices X and Y, marginals a and b, and lineage information across time points, we compute distance matrices C^X^, C^Y^ and C, and use moslin to reconstruct a coupling matrix P, probabilistically matching early to late cells. The marginals may be used to quantify measurement uncertainty or cellular growth and death. The coupling matrix P may be analyzed directly or passed to CellRank^22^ to compute fate probabilities, driver genes and expression trends- or cascades. Figure created using BioRender.

We designed moslin for time-series scLT studies (Methods). These record evolving clonal relationships using a variety of approaches, including Cas9-induced scars^10–13,25^ and naturally-occurring mutations^26,27^. We refer to the entirety of any such genomic lineage information in a single cell as a “barcode” and stress that moslin is applicable to any kind of barcode information.

Applying scLT to in-vivo systems usually requires that each time point corresponds to a different individual. We relate to this experimental design as “independent clonal evolution” (ICE), as barcode generation proceeds independently in each individual. While barcodes can be directly compared within one individual to estimate lineage trees^10–13,16,18,19^, they are incompatible across different individuals and hence time points. However, gene expression continues to be comparable across time points, giving rise to a hybrid setting where we may relate lineage or gene expression within or across time points, respectively (Fig. 1b and Methods).

To link cells from an early (t_1_) to a late (t_2_) time point, we make two major assumptions: (i) cells change their molecular state gradually, and (ii) lineage distances are, on average, conserved between time points. By lineage distance, we mean the degree to which two cells have diverged on the lineage tree. We designed moslin using the flexible framework of Optimal Transport^28,29^ (OT), which allows us to include both assumptions into a single cost function (Fig. 1c, Methods, and Supplementary Note 1).

The first assumption forms the basis of many successful pseudotime algorithms^1,30–34^; we include it in moslin using a Wasserstein (W)-term, which encourages links between cells with similar gene expression. Briefly, the W-term sums over all combinations of early and late cells, aiming to find a probabilistic mapping that minimizes the overall cost of transporting cells^1^ (Methods). The second assumption implies a type of lineage concordance: cell pairs at t_1_ should be mapped to cell pairs at t_2_ with similar relative lineage distances. We include this assumption in moslin using a Gromov-Wasserstein^35^ (GW)-term (Methods and Supplementary Note 1). Briefly, the GW-term sums over all pairwise combinations of early and late cells, aiming to find a probabilistic mapping that minimizes the discrepancy between pairwise lineage distances (Methods).

We balance both terms with an αparameter between 0 and 1, corresponding to W and GW terms, respectively^36^. This parameter allows us to tune the weight given to gene expression and lineage information. Further, we add entropic regularization at weight ϵto our objective function to speed up the optimization and to improve the statistical properties of the solution^29,37,38^. Thus, moslin solves a Fused Gromov-Wasserstein^36^ (FGW) problem with hyperparameters αand ϵ(Fig. 1c, Methods, and Supplementary Note 1).

Inputs to the moslin workflow are gene expression matrices X at t_1_ and Y at t_2_, as well as lineage information (Fig. 1d and Methods). In the first step, we compute cost matrices C and C^X^, C^Y^, representing expression and lineage distances, respectively. We quantify expression distance across time points using squared Euclidean distance in a latent space^1^, computed using PCA or scVI^39^. To quantify lineage distance within each time point, we either work with Hamming distance among raw barcodes or with the shortest path distance among reconstructed lineage trees^10–13,16,18,19^ (Methods). The choice of lineage distance metric depends on the structure of the lineage information, the expressibility of the barcodes, and the quality of tree reconstruction. In a second step, moslin solves the FGW problem to find an optimal coupling matrix P, relating cells at t_1_ and t_2_. The coupling simultaneously minimizes expression distances according to C and maximizes lineage concordance according to C^X^ and C^Y^, using the W and GW terms, respectively. For each t_1_ cell i, the vector P_i,:_ quantifies lineage and state-informed transition probabilities towards any t_2_ cell j. Finally, we use the coupling matrix P to compute ancestor and descendant probabilities^1^ directly in moslin and pass it to CellRank^22^ for further analysis.

Following previous successful approaches that link cells across time points using OT^1,21^ or related approaches^2^, we optionally include prior information about cellular growth and death into our objective function. We accomplish this by adjusting the marginal distributions passed to moslin, such that cells likely to proliferate or die can distribute more or less probability mass, respectively (Fig. 1d). We calculate growth and death rates based on prior knowledge or curated marker gene sets^1^. Our implementation additionally includes an unbalanced formulation^29,40,41^, which accounts for uncertain growth and death rates, as well as for stochastic cell sampling (Methods).

### Moslin accurately reconstructs simulated trajectories

We assess moslin’s performance on two simulated datasets. As an initial verification, we consider simulated single-cell transcriptome trajectories using a setting suggested by Forrow et al.^21^. In this simplified setting, all meaningful dynamics occur in two dimensions, representing two genes. A biologically plausible trajectory structure is prescribed via a vector field that cells follow through diffusion and occasional cell division. A lineage barcode, including random mutations, is assigned to each cell and inherited by its descendants.

We consider four different trajectories of increasing complexity: (i) *bifurcation* (B), where a single progenitor cell type splits into two descendant cell types, (ii) *partial convergent* (PC), where two initial clusters split independently, and following the split, two of the resulting four clusters merge for a total of three clusters, (iii) *convergent* (C), where two initial clusters converge to a single final cell type, and (iv) *mismatched clusters* (MC), where two initial clusters each split into two late-time clusters and cells from two of the resulting late-time clusters are transcriptomically closer to early cells that are not their ancestors (Fig. 2a, see ref.^21^).

**Fig. 2.**
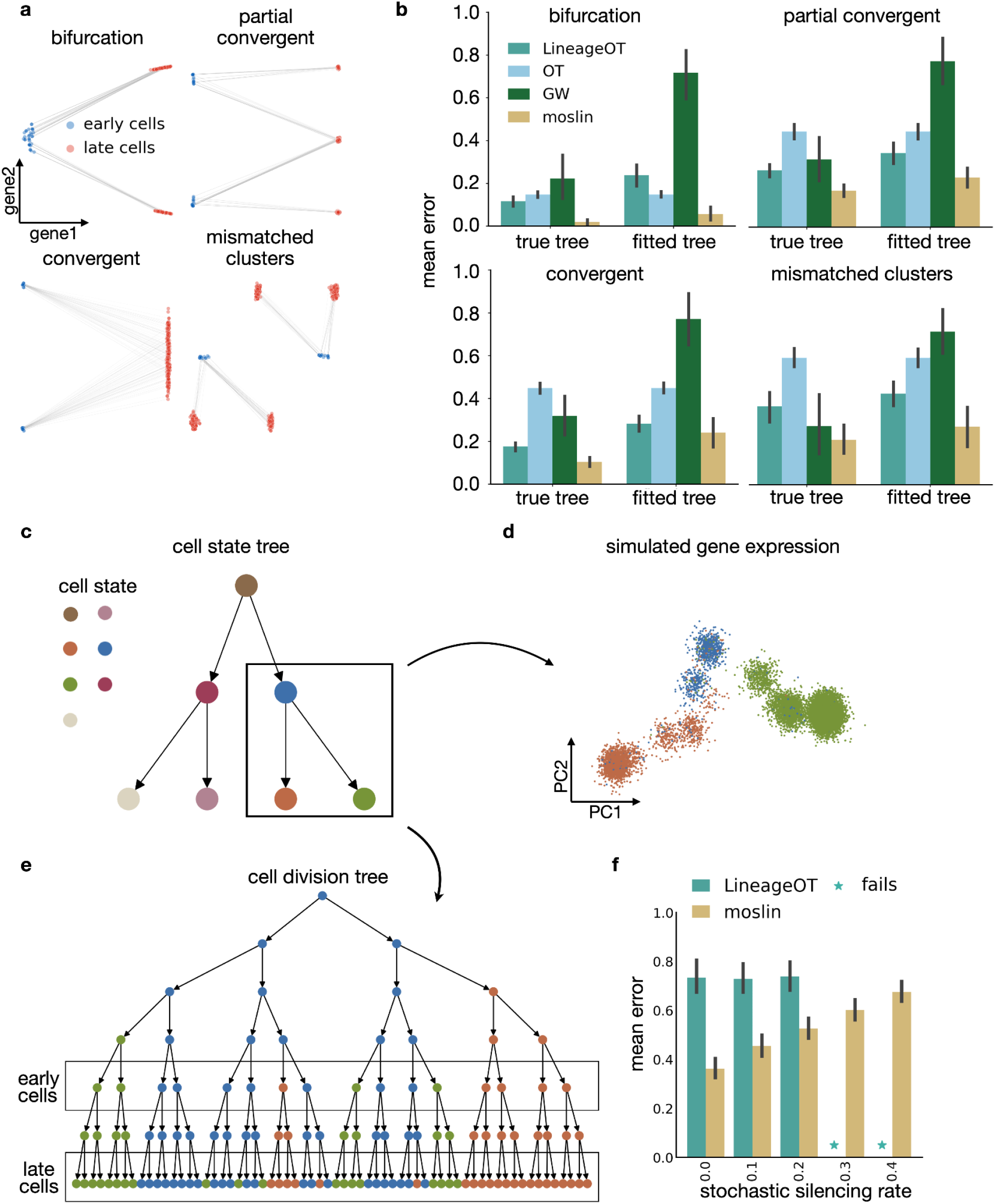
Moslin obtains accurate couplings for simple and complex trajectory topologies. **a**. Visualization of the four different kinds of simulated trajectories in gene expression space. **b**. Each subplot presents the evaluation of a different simulated trajectory. Per trajectory, the mean error (the mean value of the ancestors and descendants error) is evaluated for the true tree or a reconstructed fitted tree for all methods, LineageOT, OT, GW and moslin (Methods). Error bars depict the 95% confidence interval across 10 random simulations. **c-e**. Simulated tree and expression using TedSim^42^. The cell state tree (**c**) defines the underlying trajectories of cell differentiation. TedSim simulations yield gene expression (**d**) and a cell division tree (**e**), which represents the true lineage and barcode for each cell. **f**. Mean prediction error of moslin compared to LineageOT. As a function of the *stochastic silencing rate*. Error bars depict the 95% confidence interval across 10 random simulations.

We benchmark the performance of moslin against the only competing method, LineageOT^21^, which uses lineage information only at the later time point. We also test two extreme cases of our moslin approach: (i) using only gene expression information in a W-term (α= 0), and (ii) using only lineage information in a GW-term (α= 1) (Methods, for moslin we perform a grid search to set the interpolation parameter α). We test all methods with two types of lineage-distance computation: (i) using the ground truth tree and (ii) using a fitted tree based on the simulated barcodes (Supplementary Fig. 1a,b). We perform a grid search for each case to find the optimal hyperparameters (Methods and Supplementary Fig. 1c,d). To quantify method accuracy, we compare gene expression of predicted and ground-truth ancestors and descendants in terms of Wasserstein distance^21^ (Methods). We normalize this value by the Wasserstein distance we obtain from an uninformative coupling, given by the marginal-outer product, to obtain ancestor and descendant errors. Each value lies between 0 (ground truth) and 1 (uninformative). Finally, to obtain a single number quantifying method accuracy, we average over ancestor and descendant errors to obtain the “mean error”.

In agreement with the original publication^21^, we find that LineageOT improves over the baseline OT setting in seven of eight cases (Fig. 2b). Moslin further improves on LineageOT, with an average improvement of 10% and 12% in the mean error across all trajectories using true and fitted trees, respectively. Across all tested methods, moslin achieves the lowest mean error across all trajectory structures and distance variants. Of note, GW performs well using ground-truth tree distances, outperforming OT in three of four cases and demonstrating the value of ground-truth lineage information. However, as expected, pure GW is heavily affected by noise in tree distances and shows the largest mean error across all trajectories on more realistic, fitted tree distances.

These results demonstrate the power of the moslin approach: while pure GW is heavily affected by noisy lineage information, moslin compensates for this noise using gene expression information. Importantly, the authors of LineageOT^21^ reported that their tree reconstruction was only moderately accurate, implying that moslin outperforms the baseline OT approach in a setting reminiscent of real scLT data. Thus, the interpolation between gene expression and lineage information allows our approach to achieve excellent performance on realistically fitted tree distances (Fig. 2b).

Next, we consider a more complex simulation using TedSim^42^, which simulates cell division events from root to present-day cells. It generates two data modalities for each cell, gene expression and a lineage barcode, defining a much more complex setting than the two-dimensional regime considered above. The cell lineage tree is simulated as a binary tree that encodes cell division events, where a predefined cell state tree dictates the allowed transitions toward terminal cell states. We cut the lineage tree at an intermediate depth to simulate an early time point and use leaf nodes for the late time point (Fig. 2c-e and Methods). We map cells from the early to the late time point, providing only lineage relationships within time points to moslin and using the lineage relationships across time points to score the quality of our reconstructed mapping.

scLT datasets often suffer from barcode detection issues, and it is, therefore, crucial to assess the performance of computational pipelines on partially-detected barcodes. In our simulations, we introduce a stochastic silencing rate (*ssr*), the rate at which individual elements of the barcode remain undetected. In this example, we test an alternative to lineage tree reconstruction and directly use the scaled Hamming distance between barcodes to measure lineage distances in moslin (Methods).

We find that moslin outperforms LineageOT across our range of *ssr* values. In particular, moslin with maximal *ssr* achieves lower mean error than LineageOT on noise-free barcodes (Fig. 2f). Critically, moslin can be used robustly even for relatively high ssr, while LineageOT fails and does not provide any mapping beyond a certain threshold (*ssr* > 0.2).

### Mapping gene expression across *C. elegans* embryonic development

To showcase moslin’s performance in a realistic setting where ground truth is still available, we consider *C. elegans* embryonic development. The adult animal consists of only 959 somatic cells^43,44^, generated following a sequence of deterministic lineage decisions. This species’ ground truth lineage tree is known^44^ and available to assess moslin’s reconstruction performance. Further, this well-studied system is a good test case to validate biological insights gained by combining moslin with CellRank for fate mapping, gene dynamics, and driver gene prediction.

Previous work mapped time-series gene expression profiles of approx. 86k single cells to individual tree-nodes^7^, providing a setting where joint lineage, state and time information is available. Not all cells in this study could be mapped unambiguously. Thus, we focus on the well-annotated ABpxp lineage, which produces mostly ciliated and non-ciliated neurons, glia and excretory cells^45^. AB is one of the founding lineages of *C. elegans*; “p” (“a”) indicates the posterior (anterior) ancestor, and “x” replaces “l” (left) or “r” (right), indicating a left/right symmetry^7,45^ (Supplementary Fig. 2a). The dataset consists of 6,476 ABpxp cells across 7 time points from 170-510 min past fertilization (Fig. 3a, Supplementary Fig. 2b-d and Methods).

**Fig. 3.**
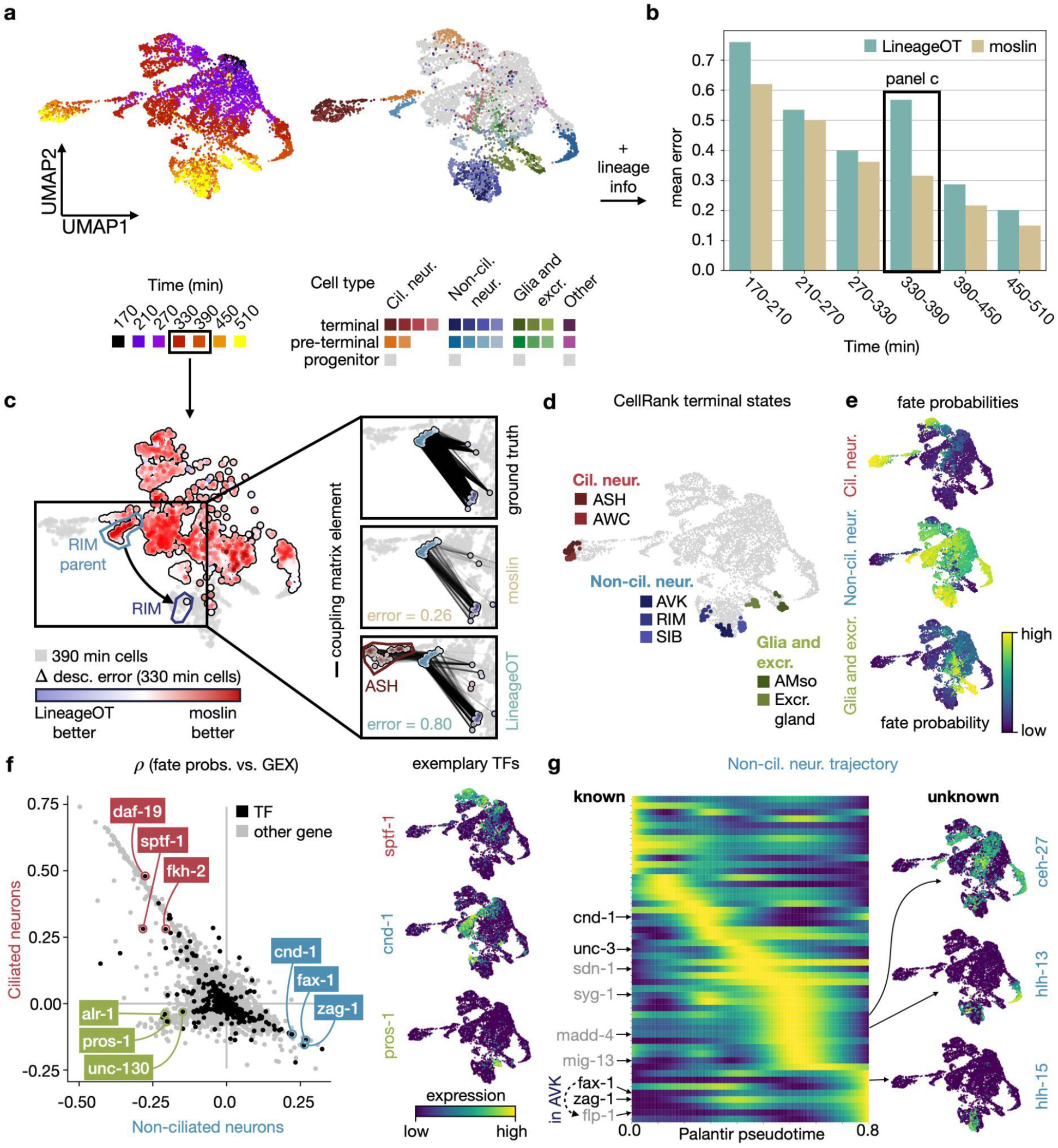
Moslin accurately captures *C. elegans* embyogenesis. **a**. UMAP^82^ of approx. 6.5k *C. elegans* ABpxp cells, colored by time point (left) and cell type (right)^7^. **b**. Bar chart of the mean error for moslin and LineageOT^21^ across time points (Methods). **c**. Left: UMAP of 330-390 min cells, colored in grey (390 min cells) or by the difference in descendant error between moslin and LineageOT (330 min cells). Black inset highlights RIM parent cells, which transition towards RIM cells^7^. Right: ground-truth, moslin and LineageOT couplings for the RIM parent population; “error” indicates the aggregated descendant error over this population (Methods) **d**. UMAP, showing the top 30 cells per moslin/CellRank^22^ computed terminal state. **e**. UMAPs of aggregated fate probabilities towards Ciliated neurons, Non-ciliated neurons, and Glia and excretory cells (Supplementary Fig. 6 and Methods). **f**. Scatter plot, showing the correlation of gene expression (GEX) with Non-ciliated-(x-axis) and Ciliated (y-axis) neuronal fate probabilities. Annotated TFs are known to be involved in the developmental trajectory they correlate with (Supplementary Table 1). Right: UMAPs, showing expression of exemplary TFs. **g**. Left: heatmap showing expression values for the top 50 predicted driver genes of Non-ciliated neurons (all gene names shown in Supplementary Fig. 10). Each row corresponds to a gene, smoothed using fate probabilities (**e**) and the Palantir pseudotime^53^ (x-axis, Supplementary Fig. 9). We annotate a few TFs, including cnd-1^48,49^, fax-1^56^, and zag-1^57–59^ (black), and other genes, including syg-1^83–85^, madd-4^86–88^, and flp-1^56,89^ (grey) that are known to be involved in the process (Supplementary Table 1). Right: UMAPs, showing expression of previously unknown predicted driver TFs.

We benchmark the performance of moslin and LineageOT across time points on the ABpxp lineage using a similar set-up as for the TedSim^42^ data, and as suggested in ref.^21^ (Fig. 2c-f). Specifically, we only provide lineage distances within time points to both methods. We compare predictions with ground-truth lineage relations across time points by calculating the mean prediction error over ancestor and descendant states (Methods). For all time point pairs, moslin outperforms LineageOT and achieves a lower mean error (Fig. 3b). These results generalize to another, distinct subset of *C. elegans* cells with precise lineage information (Supplementary Fig. 3).

The mean error difference between moslin and LineageOT is largest on the 330/390 min pair of time points. To illustrate this point, we zoom in on the difference between moslin and LineageOT per 330 min-cell (Fig. 3c and Supplementary Fig. 4). As an example, we pick a pre-terminal population of RIM (non-ciliated) neurons for which moslin’s descendant error is much smaller compared to LineageOT’s. We find that moslin correctly links these cells to RIM neurons, while LineageOT predicts many erroneous connections with ASH (ciliated) neurons (Fig. 3c and Methods).

Going beyond a single pair of time points, we combine moslin’s couplings across all time points to study *C. elegans* embryogenesis using CellRank^22^, a computational fate mapping tool. Embryogenesis, especially at later stages, is a loosely synchronized process where embryos of similar ages can represent slightly different developmental stages. This holds in particular for our *C. elegans* example, where developmental time per cell was estimated by comparing to bulk expression data^7^. Thus, we expect to find cells of slightly different maturity stages within each assigned time point. To account for developmental asynchrony, CellRank computes, for each time point, a transition matrix reflecting undirected gene expression similarity. These within-time point transition matrices are combined with moslin’s across time-point coupling matrices to yield the final transition matrix, reflecting cellular dynamics within and across time points (Methods). When we use the final transition matrix to simulate 500-step random walks from the 170 min time point, we find that these terminate in the known terminal cell types, recapitulating the established developmental hierarchy (Supplementary Fig. 5a,b).

Using this transition matrix, we set out to study gene dynamics and fate choice among ABpxp cells. As a first step, we use moslin/CellRank to compute seven terminal states and recover known Ciliated-neuronal, Non-ciliated-neuronal, Glia and excretory subtypes^7^ (Fig. 3d). The terminal states we identify are among the best-resolved cell types for Ciliated-neuronal, Non-ciliated-neuronal, Glia and excretory groups in terms of cell number (Supplementary Fig. 2d). Thus, we successfully capture representative candidates of each group. As expected, predicted terminal states mostly consist of late-stage cells, and each only contains cells from a single cell type (Supplementary Fig. 5c,d).

We aggregate the seven terminal states into three groups: Ciliated neurons, Non-ciliated neurons, and Glia and excretory cells. Next, we use CellRank to compute fate probabilities towards these groups (Fig. 3e and Supplementary Fig. 6). In agreement with known biology, moslin/CellRank predicts most progenitors in the ABpxp lineage to transition towards Non-ciliated neurons^7^ (Supplementary Fig. 7a,b). For each of the three terminal cell groups, predicted fate probabilities are significantly higher among cells from the corresponding known pre-terminal populations (Supplementary Fig. 7c and Methods). We correlate fate probabilities with gene expression to identify putative driver genes for each of the three trajectories. Focusing our attention on *C. elegans* transcription factors^46^ (TFs), we automatically recover known drivers for each trajectory, including sptf-1 for Ciliated neurons^47^, cnd-1 for Non-ciliated neurons^48,49^, and pros-1 for Glia and excretory cells^50–52^ (Fig. 3f, Supplementary Fig. 8, Supplementary Table 1 and Methods).

Finally, to study the temporal dynamics of fate decisions during *C. elegans* embryogenesis, we compute a pseudotime using Palantir^53^, starting in a 170 min-cell (Supplementary Fig. 9a). As expected, pseudotime values increase across time points (Supplementary Fig. 9b). Focusing on the Non-ciliated neuron trajectory, we compute the 50 top-correlated genes with Non-ciliated fate probabilities. For each of these genes, we combine the Palantir pseudotime with moslin/CellRank fate probabilities to compute smooth expression trends (Methods and Supplementary Table 1). Sorting expression trends by their pseudotime-peak and plotting them in a heatmap reveals a sequential activation pattern (Fig. 3g and Supplementary Fig. 10). Our results show that some TFs with known function in Non-ciliated neuron generation, including cnd-1^48,49^ or unc-3^54,55^, are activated before others, including fax-1^56^ and zag-1^57–59^ (Supplementary Table 1). In particular, our activation pattern predicts that fax-1 is activated before flp-1, a known regulatory interaction in (non-ciliated) AVK cells^56^.

While many moslin/CellRank predicted driver genes had known functions in Non-ciliated neuron generation, we also identify candidate driver genes that are novel, to the best of our knowledge. In particular, our results predict ceh-27, hlh-13 and hlh-15 as putative drivers (Fig. 3g). ceh-27 is a homeobox TF, a class of TFs known to be crucial for C. elegans neurogenesis^60,61^. While previous work^60^ reported ceh-27 expression in Non-ciliated neurons, the TF has no known function in fate specification towards these neurons. hlh-13 and hlh-15 are basic helix-loop-helix TFs; hlh-15 is known to be involved in *C. elegans* aging^62^.

### Moslin determines the dynamics of transient fibroblasts in heart regeneration

The zebrafish heart regenerates after injuries, such as ventricular resections^63^ or cryoinjuries^64–66^. A previous study used the integrated lineage-tracing and transcriptome profiling technique LINNAEUS^10^ to generate a dataset of approximately 200,000 single cells in the zebrafish heart across four time points: before injury (control), three days after injury (3dpi), seven days after injury (7dpi) and thirty days after injury (30dpi). This dataset includes inferred lineage trees and cell type annotations for each time point^67^ (Fig. 4a).

**Fig. 4.**
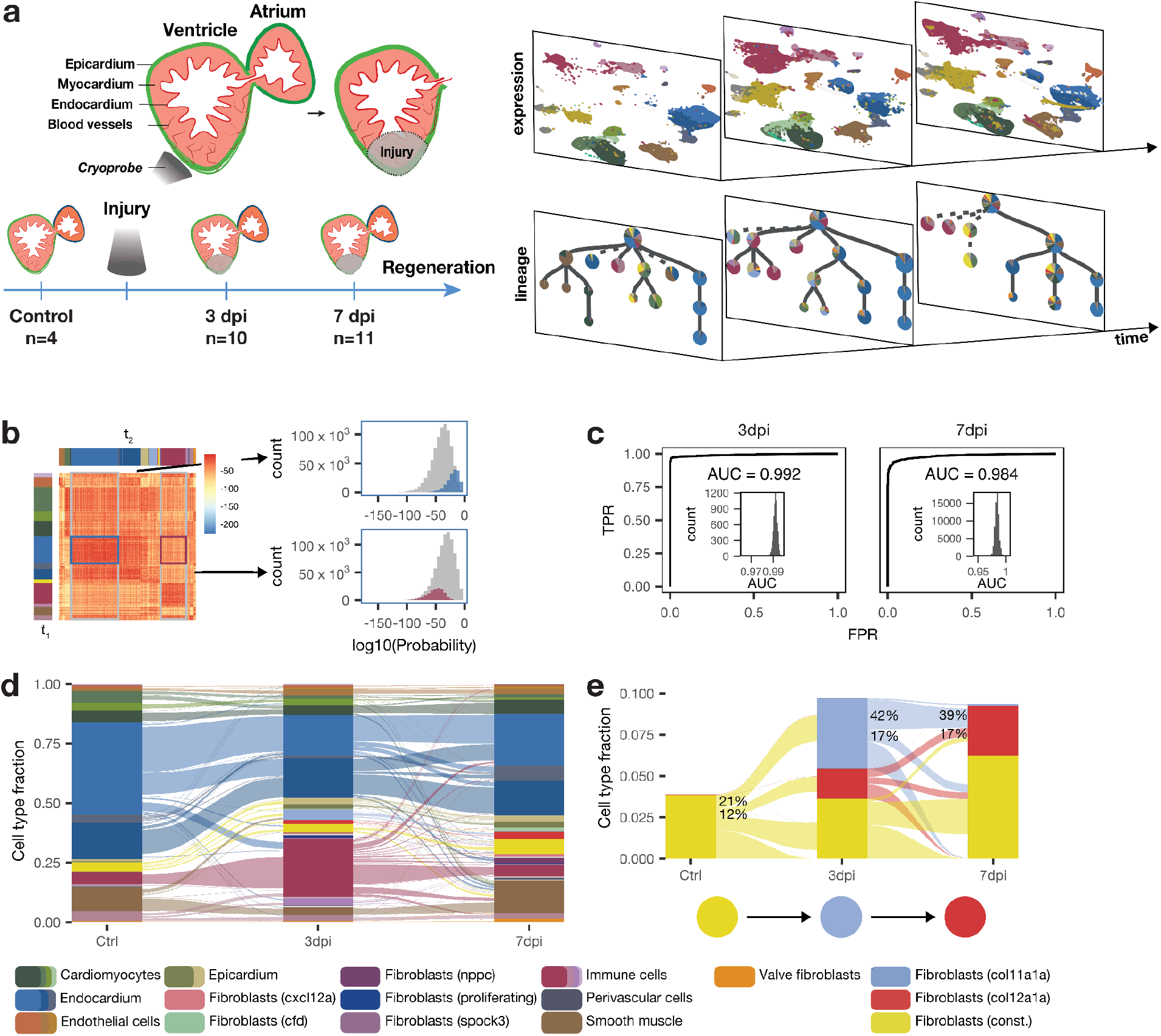
Moslin recovers lineage relations among transient fibroblast subsets. **a**. Underlying data describes zebrafish heart regeneration, measured through single-cell transcriptomic and lineage profiling before injury (n=4), at 3dpi (n=9) and 7dpi (n=7)^67^. **b**. Welch’s t-test for cell type persistence (Methods). **c**. ROC curves for same-cell type ancestors. **d**. Flow diagram of cell type transitions. **e**. Flow diagram of transient epicardial fibroblasts corroborates col11a1a fibroblasts as an intermediary state between constitutive and col12a1a fibroblasts.

One key result from this study was the emergence of several transcriptomically distinct fibroblast substates during regeneration. Analysis of lineage trees created with LINNAEUS showed that some transient states originate from the endocardial layer and others from the epicardial layer. The persistent constitutive fibroblasts share a lineage with the epicardial layer as well. One state from the epicardial layer, a fibroblast subtype characterized by a high col12a1a-expression, called col12a1a fibroblasts, was shown to be essential for regeneration: ablation of col12a1a fibroblasts strongly reduces the regenerative capacity of the zebrafish heart. Another epicardial-based transient state, the col11a1a fibroblast state, characterized by high col11a1a expression, only occurs at 3dpi, and its role is unclear. This state could lead to col12a1a fibroblasts, or it could be independent. Since the original analysis was restricted to individual time points, this question could previously not be resolved, which precluded further analysis of the underlying regulatory interactions. We reasoned that we could characterize this relationship by combining time points using moslin.

We apply moslin on all single cells in this dataset with lineage information - approximately 44,000 single cells from 20 individual animals across ctrl, 3dpi and 7dpi. We embed the transcriptomic readout of all single cells with lineage information into a joint latent space using scVI^39^, retaining the original cluster annotations. We calculate lineage distances as shortest path distances along the original reconstructed trees and use moslin to calculate couplings between cells at consecutive time points, using the interpolation parameter α= 0. 5.

Initially, we validate the performance of moslin in this challenging regeneration setting. We design a test around the assumption that most persistent cell states should be their own precursor; for example, precursors of atrial endocardial cells at 7dpi should be atrial endocardial cells at 3dpi. We use a Welch-test-based framework to test whether cells of type A at t_2_ are significantly coupled to cells of type A at t_1_, or to any other cell type B at t_2_ (Fig. 4b and Methods). Under the expectation that a significant coupling of A at t_1_ to A at t_2_ is a true positive and a significant coupling to any other cell type is a false positive, we visualize receiver operating characteristic (ROC) curves for control-3dpi and 3dpi-7dpi couplings. Areas under the ROC curve (AUCs) of 0.992 and 0.984, respectively, show that moslin can be used to determine cell state relationships across time (Fig. 4c and Methods).

In this framework, we test a cell type A at t_1_ against all cell types at t_2_; all but one of these tests (namely, the one where we test A at t_1_ against A at t_2_) is supposed to yield a negative result. To ensure AUCs are not inflated by this high amount of true negatives, we also test whether cells of type A at t_2_ are significantly coupled to cells of type A at t_1_ or to the ensemble of other cells. Here we find AUCs of 0.9999 and 1 (Supplementary Fig. 11 and Methods). Finally, we find moslin’s performance decreases by 3% (from AUC 0.99 to 0.96 at both time points) if ϵis increased to 0.1; variations in other hyperparameters yield performance changes below 1% (Supplementary Fig. 11), showing that moslin is robust to hyperparameter changes.

We next investigate the origins of transient fibroblast substates, including col11a1a and col12a1a fibroblasts. In particular, the previously published analysis had left room for two hypotheses: either col11a1a fibroblasts are an intermediary state between constitutive and col12a1a fibroblasts, or these two fibroblast states arise from constitutive fibroblasts independently. We calculate couplings with moslin, take weighted averages of cell type frequencies over separate organisms, and aggregate couplings between cell types to quantify cell type transitions during regeneration (Supplementary Fig. 12 and Methods). As expected, we observe that persistent cell types couple strongly to themselves (Fig. 4d).

Furthermore, we observe that constitutive fibroblasts preferentially generate col11a1a fibroblasts, and that most col12a1a fibroblasts originate from col11a1a fibroblasts: 21% (95% confidence interval: 15-28%) of the mass generated by constitutive fibroblasts at control goes towards col11a1a fibroblasts, whereas only 12% (95% confidence interval: 6-19%) goes directly towards col12a1a fibroblasts. At 3dpi, 42% (95% confidence interval: 22-56%) of the mass generated by col11a1a fibroblasts goes towards col12a1a fibroblasts, which constitutes 39% (95% confidence interval: 26-47%) of the col12a1a fibroblast mass at 7dpi (Fig. 4e). Confidence intervals for the frequencies and couplings were constructed by subsampling (Methods).

Taken together, this suggests that the majority of col12a1a fibroblasts is generated by constitutive fibroblasts that transition through a col11a1a-expressing state (Fig. 4e). We hypothesize that the 3dpi col12a1a fibroblasts that seem to originate directly from constitutive fibroblasts have actually transitioned through a col11a1a fibroblast state between injury and 3dpi. Our findings demonstrate the added value of temporal lineage models like moslin in analyzing scLT time-course data.

## Discussion

We demonstrate that combining intra-individual lineage similarity with inter-individual gene-expression similarity improves trajectory reconstruction substantially for in-vivo single-cell lineage-tracing (scLT) data. moslin outperforms competing methods on simulated and real data by interpolating between Wasserstein and Gromov-Wasserstein regimes and using lineage information at both time points. Crucially, we highlight in simulations that moslin compensates for noisy lineage relations through gene expression information, rendering our method suitable for real scLT data. We illustrate moslin’s capability to recover cell-state trajectories from real scLT data in zebrafish heart regeneration^67^, where we predict a new origin for regenerative activated fibroblast states. Importantly, moslin is the first computational method with demonstrated success in this challenging real data setting.

Moslin’s key advantage over previous analysis paradigms for in-vivo scLT data is that it relates cells across time points rather than focusing on individual, isolated time points. While tree reconstruction from a single time-point of lineage-traced cells can uncover shared lineage ancestry^10–13,16,18,19^, it falls short of characterising the molecular properties of these ancestors. Moslin links putative ancestors to their descendants based on lineage and gene expression information; this enables us to relate the different activated fibroblast states as a function of the time past injury, a hypothesis that remains to be validated experimentally. Cell states undergo far-reaching changes over time in many situations such as cancer, cardiovascular- and neurodegenerative diseases. To understand the gene regulatory events that underlie these changes, it is crucial to identify the corresponding sequence of state transitions. Moslin now provides a unified framework for this identification from time-resolved single-cell lineage tracing studies.

Under the hood, moslin is based on moscot, a robust and easy-to-use framework for OT applications in single-cell genomics. As such, it benefits from moscot’s interoperability with the scverse^68,69^ ecosystem and can take advantage of future moscot improvements concerning scalability and usability. Moslin’s interface with CellRank^22^ grants it access to a range of established, constantly growing downstream-analysis functions. We demonstrate the power of combining moslin with CellRank on the *C. elegans* data, where their combination reveals long-range state-change trajectories, driver genes, and temporal dynamics. Moslin’s couplings could further be employed to regularize the inference of gene regulatory networks^70,71^, or to improve perturbation predictions^72^.

In this study, we focus on the independent clonal evolution experimental design because it allows us to apply our method to in-vivo scLT data. In this setting, lineage relationships are only comparable within one time point. In contrast, for in-vitro experiments, cells from the same population can be sampled at different time points, rendering their lineage information directly compatible across time points. Previously, OT-like approaches^14,15^ have been suggested for this clonal resampling experimental design^24^. Moslin could be extended towards this setting by adjusting the cost-matrix definition.

While moslin is robust to noise in lineage information, it will benefit from improved experimental lineage tracing technologies. Recent innovations, including mitochondrial lineage tracing^26,27,73^ and base/prime editing^74–77^, represent compelling use cases for moslin. Improved lineage resolution will allow our method to yield highly-accurate trajectory reconstructions in challenging disease contexts like cancer or inflammation.

Currently, moslin is limited to one replicate per time point. In the zebrafish data^67^, where several replicates per time-point are available, we address this by computing pairwise replicate linkages across time points and aggregating our insights across these. With the increasing popularity of scLT approaches, we expect more complex, multi-replicate time series to become available. For these, as an alternative to the aggregation approach above, we envisage a two-step process, first computing a consensus lineage representation per time point across replicates, and second, linking the consensus representations across time points.

Moslin could further be extended towards multi-modal scLT data^78,79^ to link molecular layers across time. For example, this could reveal how epigenetic changes manifest in altered gene expression dynamics^80,81^. Additionally, spatially-resolved lineage tracing data would enable moslin to regularise the coupling computation further using spatial neighbourhoods. In this setting, moslin’s inferred trajectories could be used to interrogate the relative contribution of internal state versus external signals towards observed fate decisions. scLT is a fast-moving field; we anticipate computational tools like moslin to play a crucial role in analyzing and interpreting novel lineage-traced datasets.

## Acknowledgements

We would like to thank Adan Forrow for helpful discussions about LineageOT, Xinhai Pan as well as Xiuwei Zhang for advice regarding their simulation tool, TedSim, Shou-Wen Wang for clarifications regarding Cospar, and Marco Cuturi for input concerning optimal transport. We would further like to thank Matthew G. Jones and all members of Nitzan and Theis labs for the great discussions. This work was supported by the BMBF (grant No. 01IS18036B and grant No. 01IS18053A), by the Helmholtz Association (Incubator grant sparse2big, grant No. ZT-I-0007), by the European Union (M.N.: ERC, DecodeSC, 101040660; J.P.J.: ERC, 715361; F.J.T.: ERC, DeepCell, 101054957), by the Wellcome Trust, Wellcome Leap, Delta Tissue [9E8E84F7-8991-4D4A-A9EC], by the DZHK (German Centre for Cardiovascular Research), by the Israel Science Foundation (grant No. 1079/21), and by the BMBF-funded de.NBI Cloud within the German Network for Bioinformatics Infrastructure (de.NBI) (031A532B, 031A533A, 031A533B, 031A534A, 031A535A, 031A537A, 031A537B, 031A537C, 031A537D, 031A538A). M. Lange further acknowledges financial support by the DFG through the Graduate School of QBM (GSC 1006) and by the Joachim Herz Foundation. Z.P is supported by a scholarship for outstanding doctoral students in data-science by the Israeli council for higher education and the Clore Scholarship for Ph.D students. M.N. is supported by an Early Career Faculty Fellowship by the Azrieli Foundation. For all support coming via EU funding, the views and opinions expressed are those of the author(s) only and do not necessarily reflect those of the European Union or the European Research Council Executive Agency. Neither the European Union nor the granting authority can be held responsible for them.

## Author contributions

M.L. conceived the project, guided by F.J.T. and M.N. J.P.J., F.J.T. and M.N. supervised the research. M.L. designed the algorithm with contributions by Z.P. and M.K., and guided by M.N. M.K. implemented the algorithm, with contributions by D.K. M.K. and Z.P. benchmarked the method across datasets. Z.P. conducted the simulations studies, with contributions by M.K. M.L. analyzed the C. elegans data, with contributions by M.K. B.S. analyzed the Zebrafish data, with contributions by Z.P. M.L., Z.P., B.S., F.J.T. and M.N. wrote the manuscript. All authors read and approved the final manuscript.

## Data availability

Raw published data for the C. elegans^7^ and Zebrafish^67^ examples are available from the Gene Expression Omnibus under accession codes GSE126954 and GSE159032, respectively. Processed data is available from figshare under https://doi.org/10.6084/m9.figshare.c.6533377.v1

## Code availability

The moslin software can be accessed via https://github.com/theislab/moslin, including documentation, tutorials and examples. Jupyter notebooks and Python scripts to reproduce our results are available via the same GitHub repository.

## Supplementary materials

Supplementary Figures: Supplementary Figures 1 - 12

Supplementary Tables: Supplementary Table 1

Supplementary Notes: Supplementary Note 1

## 1 The moslin algorithm

### 1.1 Introduction and model overview

Moslin is an algorithm aimed at linking single-cell profiles across experimental time points. Computational linkage is required as sequencing is destructive; moslin thus allows linking molecular differences among cells at early time points with their eventual fate outcome at later time points. Critically, moslin uses molecular similarities and lineage tracing information to solve this challenging reconstruction problem. Specifically, moslin is applicable to dynamic, CRISPR-Cas based approaches^1–12^ that record lineage relationships in vivo. While previous analysis approaches for this type of lineage tracing data remained limited to individual, isolated time-points ^4,5,13–21^, moslin embeds clonal dynamics in their temporal context.

#### Moslin’s inputs

The input to moslin are pairs of state matrices and linege information (*X* ∈ ℝ ^*N×G*^, *ξ*) and (*Y* ∈ ℝ ^*M×G*^, *ζ*) corresponding to *N* and *M* observed cells at early (*t*_1_) and late (*t*_2_) time points. State matrices *X* and *Y* typically represent gene expression (scRNA-seq) across *G* genes; however, moslin can also be applied to modalities like chromatin accessibility. The lineage information arrays *ξ* and *ζ* contain the lineage tracing outcome for every cell; their exact nature depends on the lineage tracing technology (Section 1.2). Optionally, moslin takes marginal distributions ***a*** ∈ *Δ*_*N*_ and ***b*** ∈ *Δ*_*M*_ over cells at *t*_1_ and *t*_2_ for probability simplex 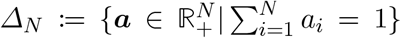. These marginals can represent any cell-level prior information; we use them to incorporate the effects of cellular growth and death.

#### Moslin’s outputs

The output of moslin is a coupling matrix *P* ∈ *U* (***a, b***) where *U* (***a, b***) is the set of feasible coupling matrices given by

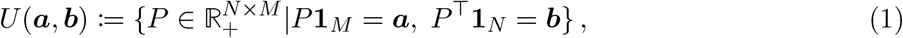

for constant one vector **1**_*N*_ = [1, …, 1]^⊤^ ∈ ℝ^*N*^. The coupling matrix *P* links cells at *t*_1_ with cells at *t*_2_; the *i*-th row *P*_*i*,:_ tells us how cell *i* from *t*_1_ distributes its probability mass across cells at *t*_2_ and the *j*-th column *P*_:,*j*_ tells us how much probability mass cell *j* at *t*_2_ receives from cells at *t*_1_. The set *U* (***a, b***) contains all matrices *P* which are compatible with the prescribed marginals ***a*** at *t*_1_ and ***b*** at *t*_2_.

With these definitions at hand, we can formalize the aim of moslin: we seek to find the coupling matrix *P* ∈ *U* (***a, b***) which simultaneously minimizes the distance cells have to travel in phenotypic space between *t*_1_ and *t*_2_ while respecting lineage relationships. We explain how we find such a matrix in Subsection 1.3

### 1.2 In vivo single-cell lineage tracing (scLT)

Moslin uses lineage tracing data to guide the reconstruction of a coupling matrix *P* between *t*_1_ and *t*_2_ cells. Early methods for lineage tracing were labor-intensive, limited to transparent organisms, and relied on manual observation of individual cells in time-lapse microscopy ^22,23^, recent approaches are sequencing-based and use heritable genetic barcodes^23–27^. While a multitude of such techniques exists, moslin is geared towards those that achieve single-cell resolution, yield joint lineage and gene expression readout, and can be applied in vivo.

#### Clonal resampling (CR) versus independent clonal evolution (ICE)

Critically, moslin is able to describe non-steady state biological processes like development or regeneration that require timeseries experimental designs to capture cell-state trajectories. Experimentally, this can be achieved using either *clonal resampling* (CR) or *independent clonal evolution* (ICE) designs, which assay cells from the same or different clones across several time points, respectively.

In clonal resampling (CR), the aim is to observe the same clone (cells sharing the same barcode) across several time points, i.e., for a single phylogenetic tree, we aim to observe some ancestral nodes besides the leaf nodes. As this approach relies on the repeated sampling of clonally related cells, it applies primarily to in-vitro settings^28–30^, in vivo transplantation settings ^28^ or in vivo regenerative systems like human PBMC and CD34+ samples ^31,32^ or the zebrafish fin ^2^. Beyond these transplantation and regenerative settings, applying time-series scLT in vivo requires independent clonal evolution (ICE), i.e., different individuals, sequenced at different time points with independent clonal evolution proceeding in each animal. This represents an additional challenge since the lineage of cells in different individuals cannot be compared directly. We designed moslin for the challenging ICE setting that allows us to model in-vivo systems.

### 1.3 Moslin’s objective function for in-vivo ICE

With the definition of ICE at hand, we return to moslin’s key task: finding a coupling matrix *P* ∈ *U* (***a, b***) which simultaneously minimizes the distance cells have to travel in phenotypic space while respecting lineage relationships. Mathematically, we cast this task as an Optimal Transport (OT) problem ^33^; in particular, we use a *Fused Gromov Wasserstein* ^34^ (FGW) formulation which allows us to include terms for across- and within time-point similarity (Supplementary Note 1). Previous single-cell methods successfully used OT to map cells across time points without lineage information^35,36^, impute gene expression in spatial data ^37^, predict perturbation response^38–40^, learn patient manifolds^41,42^, integrate data across modalities^43^ and infer cell-cell communication ^44^. In particular, we make the following assumptions (A):

- A1: cells change their state gradually; overall, they minimize the distance traveled in phenotypic space between *t*_1_ and *t*_2_.
- A2: on average, molecular similarity is conserved between *t*_1_ and *t*_2_; similar cell pairs at *t*_1_ are likely to transition into similar cell pairs at *t*_2_.
- A3: on average, lineage relations are concordant across time-points; cells with similar lineage history at *t*_1_ are likely to transition into cells with similar lineage history at *t*_2_.

All three assumptions may be challenged in practice:

- Batch effects and incomplete molecular information challenge A1.
- Rapid transcriptional convergence and divergence challenges A2.
- Noisy or incomplete lineage readout challenges A3.

Thus, rather than enforcing A1-A3 exactly, we design custom cost functions to balance them in our FGW objective function; individual cells may violate any combination of assumptions at the cost of incurring a penalty.

#### A combined approach for in vivo scLT data

In ICE, gene expression information is comparable across time points but lineage information is not (Section 1.2). Our FGW setting allows us to define terms that handle both type of information:

- A linear *Wasserstein* (W) term for comparable features, encouraging A1. This term quantities gene expression similarity.
- A quadratic *Gromov-Wasserstein* (GW) term for incomparable features, encouraging A2 and A3. This term quantifies lineage and expression concordance.

#### The W term for individual comparisons

To encourage A1, we consider a W term ^33^ which compares individual cells in the source (*t*_1_) and target (*t*_2_) distributions in terms of their gene expression vectors. Given gene expression vectors (***x***_*i*_, ***y***_*j*_) ∈ 𝒳 × 𝒴, we construct a cost matrix, 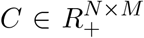 with *C*_*ij*_ = *c*(***x***_*i*_, ***y***_*j*_) for cost function *c*. An entry in the cost matrix, *C*_*ij*_, depicts the distance between cells *i* and *j* according to the cost function *c*. We define the cost function to represent squared euclidean distance in a joint latent space over *X* and *Y*, computed using PCA or scVI ^45^. Formally, the mapping problem is defined as

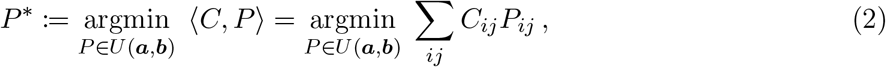

for optimal coupling matrix *P*^*^. This objective function defines a convex linear program; the optimal *P*^*^ will be the one accumulating the lowest cost according to *C* when transporting cells from *t*_1_ to *t*_2_.

#### The GW term for pairwise comparisons

To encourage A2 and A3, we consider a GW term ^33,46,47^ which compares cell pairs in the source (*t*_1_) and target (*t*_2_) distributions in terms of their gene expression and lineage information. Given gene expression vectors and lineage information, we define two independent cost matrices, 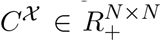 and 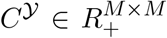 with 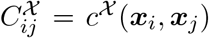 and 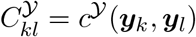 for cost functions *c*^𝒳^ and *c*^𝒴^.

Focusing on the early time point, consider latent space samples ***x***_*i*_ and lineage information *ξ*_*i*_. Define the composite *t*_1_-cost function

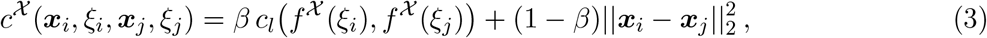

for parameter *β* ∈ [0, 1], controlling the weight given to lineage versus molecular state, mapping function *f* ^𝒳^, providing a representation of the lineage information at *t*_1_, and lineage distance function *c*_*l*_. Lineage information is typically noisy and incomplete; we include molecular similarity at weight (1 − *β*) as a regularization. Moslin supports two ways of representing lineage information:

- barcode representation: *f*^𝒳^ is the identity and *c*_*l*_ quantifies hamming distance between raw barcodes.
- lineage tree representation: *f* ^𝒳^ is a lineage-tree reconstruction computed using a method like Cassiopeia ^13^ or LINNAEUS ^1^ and *c*_*l*_ quantifies shortest path distance along reconstructed trees.

We employ an analogous set of definitions for the *t*_2_-cost function *c*^𝒴^. We apply these cost functions to all (pairs of) cells to yield the cost matrices *C*^𝒳^ ∈ ℝ^*N×N*^ and *C*^𝒴^ ∈ ℝ^*M×M*^. With the cost matrices at hand, we define a quadratic GW term that compares pairwise distances across time-points,

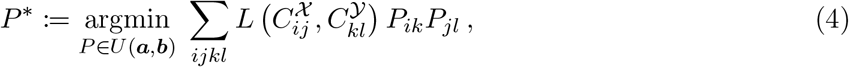

for some distance metric *L* that compares cost-matrix entries. By default, we use the *l*_2_ distance in moslin. Intuitively, this term encourages similar cells at *t*_1_ to be matched to similar cells at *t*_2_.

#### Moslin’s Fused Gromov-Wasserstein (FGW) approach

To simultaneously encourage A1, A2, and A3, we combine the W with the GW term to yield moslin’s objective function for in-vivo ICE data,

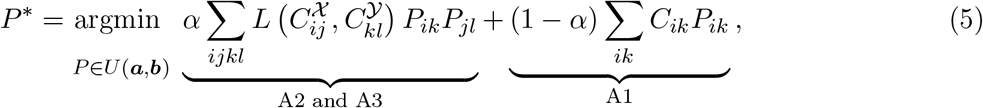

which is known as a *Fused Gromov-Wasserstein* (FGW) problem ^34^ (Supplementary Note 1). The parameter *α* ∈ [0, 1] controls the interpolation between the W and GW terms. Using this interpolation, we jointly optimize the coupling with respect to gene expression and lineage information.

#### Entropic regularization and optimization

The combined objective of Equation (5) defines a quadratic programming problem; to introduce a notion of uncertainty and to speed up the optimization, we follow previous approaches ^35,48^ and include and entropy regularization term,

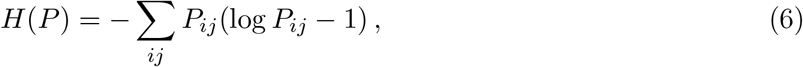

and the regularized FGW objective reads

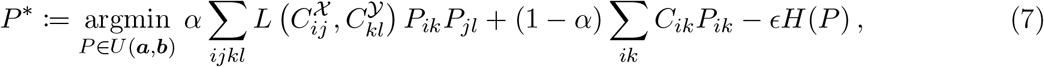

for regularization strength *ϵ*. Intuitively, the entropy term *H*(*P*) favors probabilistic over deterministic couplings. We optimize the entropy-regularized FGW objective function using a mirror descent scheme; each inner iteration of the algorithm reduces to well-known Sinkhorn iterations^33,48^ (Supplementary Note 1). To determine convergence, we check whether the current and previous regularized OT costs are close using `jax.numpy.isclose(…, rtol=R_TOL), with R_TOL = 1e-3` by default.

#### Marginals endcode prior biological information

If additional information about sampled cells is available, e.g., growth- and death-rates, uncertainty, etc., we incorporate them via the marginals ***a*** and ***b***. If no additional information is available, we assign them uniformly. By default, in moslin, we choose the right marginal ***b*** uniformly, *b*_*j*_ = 1*/M* ∀*j* ∈ {1, …, *M*}, and adjust the left marginal to accommodate cellular growth and death between *t*_1_ and *t*_2_,

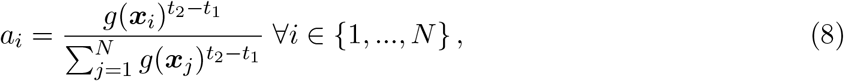

where *g* : ℝ^*D*^ → R is modeled as the expected value of a birth-death process with proliferation at rate *β*(***x***) and death at rate *δ*(***x***), thus *g*(***x***) = *e*^*β*(***x***)−*δ*(***x***)^ for *β*(***x***) and *δ*(***x***) estimated from curated marker gene sets for proliferation and apoptosis, respectively^35^.

#### Accommodating uncertainty in the inputs

As we estimate growth- and death rates from marker genes, they represent a noisy estimate of the underlying ground truth growth- and death rates. In addition, we randomly sample cells from a population, which leads to deviations from the groundtruth cell-type proportions.

Accordingly, we allow small deviations from the exact marginals ***a*** and ***b*** in an unbalanced FGW framework ^49^ where we replace the hard constraint *P* ∈ *U* (***a, b***) with soft Kullback–Leibler (KL) divergence penalties, giving rise to moslin’s final objective function for time-series scLT data. To control the weight given to left (***a***) and right (***b***) marginal constraints, we use two parameters *τ*_*a*_, *τ*_*b*_ ∈ (0, 1) (Supplementary Note 1). For the optimization, we employ the algorithm presented by Séjourné et al. ^49^ which is based on a bi-convex relaxation leading to alternate Sinkhorn iterations.

#### Implementation

Moslin is available at https://github.com/theislab/moslin. Under the hood, moslin is based on moscot, our open-source framework for **M**ulti-**O**mic **S**ingle-**C**ell **O**ptimal **T**ranport. moscot is a scalable, easy-to-use, open-source solution for OT-based analysis in single-cell genomics; it interfaces with optimal transport tools ^50^ (OTT) in the backend to support GPU acceleration and just-in-time compilation via JAX^51^.

### 1.4 Downstream usage of coupling matrices

Once we have identified the optimal coupling matrix *P*, we use it to link observed cells between *t*_1_ and *t*_2_. Note that the coupling matrix *P* combines the information from molecular similarity and lineage history; thus, all downstream analysis is lineage- and state informed.

Consider a *t*_1_ cell state 𝒫 of interest. This state could represent, e.g., a rare or transient population with unknown position in the differentiation hierarchy. Define the corresponding normalized indicator vector,

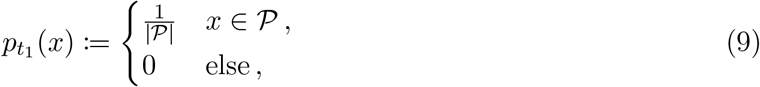

where *x* is a cell from *t*_1_ and |𝒫| corresponds to the number of cells in state 𝒫. Following Schiebinger et al. ^35^, we compute *t*_2_ descendants of cell state 𝒫 by a push-forward operation of 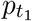,

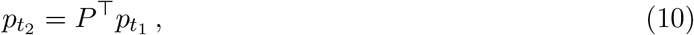

where 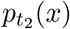 is the probability mass that cell state 𝒫 distributes to cell *x* at *t*_2_. Similarly, to compute ancestors of a cell state 𝒬 at *t*_2_, consider the corresponding normalized indicator vector 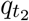. To compute the ancestor distribution, we use a pull-back operation,

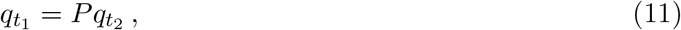

where 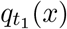 is the probability mass that cell *x* contributes towards cell state 𝒬 at *t*_2_. For further downstream analysis, e.g. to identify initial and terminal states, driver genes of fate decisions, and gene expression trends, we interface with CellRank ^52^, a fate mapping toolkit which analyzes our coupling matrices using a Markov framework.

#### Coupling cells across more than two-time points

Moslin relates cells across more than two-time points; consider a time-series experiment with sequencing at time points {*t*_1_, …, *t*_*T*_}. Following Schiebinger et al. ^35^, we solve for individual pairwise couplings between adjacent time points; this yields coupling matrices 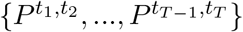. We construct longer-range couplings by matrix-multiplying individual couplings. For example, to couple initial-day cells to final-day cells, we obtain

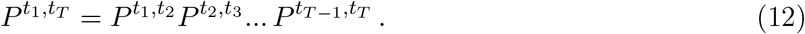

We compute ancestors and descendants for multi-day couplings in the same way as above (Equations (10) and (11)).

## 2 Datasets

### 2.1 2-gene simulations

We use a simulation setting suggested by Forrow and Schiebinger 53 which constructs a vector field to recreate a biologically plausible trajectory structure. Under the simulation, cells follow the vector field with diffusion and occasional cell division. The simulation assigns a heritable lineage barcode that is randomly mutated, to each cell. Four different types of trajectories, of increasing complexity, are considered in this simulated setting:

1. bifurcation (B): a simple bifurcation of a single progenitor cell type into two descendant cell types.
2. partial convergent (PC): two initial clusters split independently, following the split, two of the resulting four clusters merge together for a total of three clusters.
3. convergent (C): two initial clusters converge to a single final cell type.
4. mismatched clusters (MC): two initial clusters both split into two late-time clusters, and cells from two of the resulting clusters are transcriptomically closer to early cells that are not their ancestors

The simulated data provides us with what Forrow and Schiebinger ^53^ define as an *embedded lineage tree*, referring to the collection of branching paths due to cell divisions within a population (whereas a lineage tree denotes the coordinate-free tree structure). For each of the trajectories, we simulate 10 different data sets with a different random seed and measure the *embedded lineage tree* at two time points (with 64 and 1024 cells respectively). All simulations were performed using the default settings provided in the LineageOT code package: https://github.com/aforr/LineageOT.

Given the simulated data, which consists of gene expression, barcodes, and the true lineage tree, we compute couplings between time points in two manners, considering the *true tree* or a *fitted tree*. For the latter, the tree is inferred using the neighbor-joining algorithm^54^ as implemented in LineageOT ^53^. LineageOT uses the tree (true or fitted) directly to compute the couplings. In moslin, we construct the lineage costs by computing distances between cells along the tree. The distance is defined as the length of a weighted shortest path found using Dijkstra’s algorithm^55^ with weights associated to edges according to “time” between two nodes. We compare the performance of moslin to LineageOT, and two extreme cases of the moslin formulation: using only gene expression in a W-term (*α* = 0), and using only lineage information in a GW-term (*α* = 1). We quantify method performance using the ancestor and descendant errors introduced in Forrow and Schiebinger ^53^. For ground truth coupling *P*^*^ and predicted coupling *P*, we compare their predicted ancestors and descendants per cell using a Wasserstein-2 distance (Supplementary Note 1). To obtain the descendant error *E*_*D*_(*P*), we compute

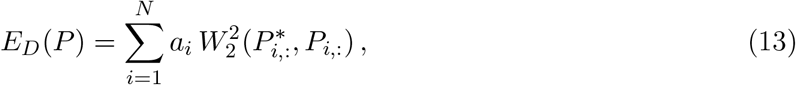

for squared Wasserstein-2 distance 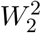 (Supplementary Note 1) and right marginal *a*_*i*_ = Σ_*j*_ *P*_*ij*_. Similarly, to obtain the ancestor error *E*_*A*_(*P*), we compute

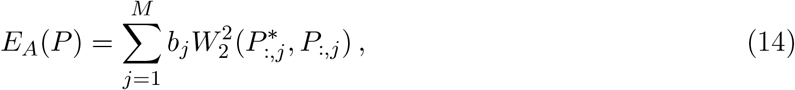

for left marginal *b*_*j*_ =Σ _*i*_ *P*_*ij*_. Note that we compare rows for *E*_*D*_(*P*) and columns for *E*_*A*_(*P*), scaled by the corresponding marginal to adapt the weight we give to each cell. Thus, a value of zero in either metric means that we are on par with the ground-truth coupling. Additionally, we independently normalize ancestor and descendant errors using the outer product of the marginals, 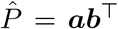, corresponding to an uninformative coupling with the same marginals as the predicted coupling *P*. Specifically, we compute 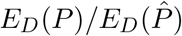 and 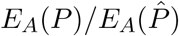, such that a value of one corresponds to an uninformative result. Our final error metric is given by the mean of the two quantities ^53^.

We perform a grid search to find the optimal parameters for each data set and method. For all settings, the entropy parameter is optimized over 15 values of *ϵ* log-spaced between 1*e* − 4 and 1*e* + 1. For moslin, we also perform a grid search for the interpolation parameter, *α* ∈ {0.1, 0.2, 0.3, 0.4, 0.5, 0.7, 0.8, 0.9, 0.95, 0.98, 0.999}.

### 2.2 TedSim simulated data

We utilize TedSim ^56^ (single-cell temporal dynamics simulator), which simulates cell division events from root cells to present-day cells, simultaneously generating two data modalities for each cell, gene expression, and a lineage barcode. The cell lineage tree is simulated as a binary tree that models the cell division events. In order to simulate diverse cell types, the notion of asymmetric divisions ^57–59^ is used. The asymmetric divisions allow cells to divide into cells with different cellular fates. One cell evolves into a new state and the other preserves the ancestor state. The evolution of cells is governed by a *cell state tree*. Two user-defined parameters control this simulation process:

1. *step*_*size*: defines the distance between two adjacent sampled states on the cell state tree. Larger *step*_*size* implies more distinct cell states along the tree.
2. *p*_*a*_: the probability for a division in the sampled tree to be asymmetric. Larger *p*_*a*_ implies rapid transitions in the sampled tree.

In accordance with the original publication ^56^, we noticed that these parameters have a small effect on the mapping accuracy hence report results for *p*_*a*_ = 0.4 and *step*_*size* = 0.4.

For the lineage information, barcodes are simulated as an accumulation of CRISPR/Cas9-induced scars along the paths from the root to all the leaf cells. Here, we add to the TedSim simulated barcodes a stochastic silencing rate, corresponding to the rate at which entire segments are removed from the barcode. With this, we aim to simulate the expected dropout due to low sensitivity of assays.

To obtain the datasets we follow the TedSim published tutorial, Simulate-data-multi.Rmd. Setting *p*_*a*_ = 0.4 and *step*_*size* = 0.4 and creating 10 different data sets using different random seeds.

Given the simulated gene expression and barcodes, we define moslin’s lineage costs as the scaled hamming distance between the barcodes, as defined by Forrow and Schiebinger ^53^. The scaling is defined such that: (i) the number of sites where both cells were measured is taken into account, (ii) the distance between two scars is twice the distance from scarred to unscarred sites. For LineageOT, similarly to the previous setting, the barcodes are used internally to construct a fitted tree. To benchmark moslin, we ran a grid search over *α* ∈ {0.1, 0.25, 0.5, 0.75, 0.9, 1} and *ϵ* ∈ {1*e* − 3, 1*e* − 4}. For LineageOT, we tested with *ϵ* ∈ {1*e* − 1, 1}.

### 2.3 *C. elegans* embryonic development

The *C. elegans* development dataset^60^ contains gene expression for approx. 86k single cells, sequenced using 10x genomics. The original authors ^60^ mapped these cells towards the known *C. elegans* lineage tree ^22^ and obtained lineage information for a subset of cells. Additionally, they mapped their data towards a bulk time-series dataset ^61^ to estimate the developmental stage of individual cells. Binning these estimated cell times gave rise to several pseudo-experimental time points, spanning 150-580 min past fertilization.

#### Preprocessing

To evaluate moslin’s performance, we required ground-truth lineage information. The original study’s ^60^ mapping inferred partial lineage information for a subset of approx. 46k cells. To obtain precise lineage information, we implemented two suggestions by Forrow and Schiebinger ^53^ :

1. Strategy 1: subsetting to the ABpxp lineage. This is a symmetric lineage where “x” indicates either the right (*“*r”) or the left (“l”) cell.
2. Strategy 2: subsetting to all cells with precise lineage information.

As the lineage for cells obtained from strategy 1 is not fully specified due to “x”, the two strategies lead to disjoint subsets of cells, allowing us to test moslin’s performance in two different scenarios.

For either cell subset, we preprocessed the data using SCANPY^62^, and used default parameters if not indicated otherwise. In particular, we normalized total counts, log-transformed the data, annotated the top 3k highly variable genes using the “seurat” flavor ^63^, and computed 50 principal components in the space of highly variable genes. To have a sufficient number of cells per time point, we removed time points that contained less than 100 cells. This left us with the following 7 time points: 170, 210, 270, 330, 390, 450 and 510 min past fertilization.

#### Embedding and cell-type labels

Using the top 10 principal components, we computed a k-nearest neighbor (kNN) graph for 30 nearest neighbors and visualize it by computing a UMAP embedding ^64^. To reduce complexity and focus on the main groups of terminal cell states, we aggregated original cluster annotations ^60^ slightly to arrive at the annotations we show in Fig. 2 and Supplementary Fig. 2. Our aggregation entailed the following steps:

- Summarize AIM, AIY, AVB, DB, PVP, RIB, RIC, SIA and RIV as “other terminal non-ciliated neurons”.
- Summarize Neuroblast_PVC_LUA and Parents_of_U_F_B_DVA as “other pre-terminal non-ciliated neurons”.
- Summarize pm7, DVA, GLR, DA and Pharyngeal_neuron as “other terminal cells”.
- Summarize AIN_parent, M1_parent, PVQ_parent, RME_LR_parent, Parents_of_Y_DA6_DA7_DA9, Parent_of_tail_spike_and_hyp10 and Parents_of_PHsh_hyp8_hyp9 as “other pre-terminal cells”.

The vast majority of cells we labeled “other terminal cells” are Pharyngeal neurons (24/30 cells), and the vast majority of cells we labeled “other pre-terminal cells” are pre-terminal hypodermis cells (Parent_of_tail_spike_and_hyp10 with 53/90 cells and Parents_of_PHsh_hyp8_hyp9 with 25/90 cells). We show the original cluster annotations, prior to aggregation, in Supplementary Fig. 2.

We labeled cells that had neither terminal or pre-terminal cell-type label (but lineage annotation) as “progenitors”. These correspond to earlier cells in the lineage tree, for which terminal identity has not been established yet.

#### 2.3.1 Benchmarking moslin with LineageOT

##### Shared moslin/LineageOT parameters and settings

We benchmarked moslin with LineageOT on the two cell subsets (Strategy 1 and 2), using the pre-processing described above. We use the marginals ***a*** and ***b*** to capture the effects of cellular growth and death, and calculate them using the lineage tree following Forrow and Schiebinger ^53^. Gene-expression distances among cells from different time-points were measured using squared Euclidean distance in the PCA space, and passed to both methods in the mean-scaled cost matrix *C*.

##### Additional moslin parameters

We did not allow for deviations from the marginals via unbalancedness in this application, as the marginals are lineage-informed and thus more accurate compared to other applications. We set *β* = 0, i.e. the GW term corresponds to pure lineage information. To construct the lineage cost matrices *C*^*X*^ and *C*^*Y*^, we compute distances between same-time point cells along the lineage tree. The distance is defined as the length of a weighted shortest path found using Dijkstra’s algorithm ^55^. The weights represent the temporal difference between a node and its parent. Additionally, we mean-scaled the *C*^*X*^ and *C*^*Y*^ cost matrices.

##### Additional LineageOT parameters

We run LineageOT following the original authors’ reproducibility repository. LineageOT runs the Sinkhorn algorithm as implemented in python optimal transport (POT)^65^ under the hood; their convergence criterion checks that the constraints imposed by the marginal distributions are satisfied within a certain threshold. We set this threshold to 10^−3^.

##### Grid search

To identify the best hyperparameters for either method per time-point pair, we run a grid search over the following parameter grid:

- Moslin:
  - *α* ∈ [0.01, 0.1, 0.25, 0.5, 0.75, 0.9, 0.95, 0.98]
  - *ϵ* ∈ [0.001, 0.01, 0.05, 0.1, 0.5]

- LineageOT:
  - *ϵ* ∈ [0.001, 0.01, 0.05, 0.1, 0.5]

For each method, the performance we report corresponds to the best performance found across this grid.

##### Mean error computation

To quantify method performance per time-point, we computed the ancestor and descendant errors over the PCA space, as described above for our simulation study. We used the mean over ancestor and descendant errors as our final accuracy metric.

##### Zoom in to the 330/390 min time point pair

To visualize the transitions predicted by moslin and LineageOT for the RIM_parent population, we selected 330 min RIM_parent cells. Out of these, we further restricted our attention to those cells assigned to the ABpxppaapa lineage; these cells represented the vast majority (80/85) of the RIM_parent population. We considered the corresponding rows in the moslin/LineageOT-predicted coupling matrices. To focus on the most confident predicted links, we only retained matrix elements exceeding 10% of the maximum coupling value, i.e. we required *P*_*ij*_ *>* 0.1 max_*ij*_ *P*_*ij*_, separately for moslin, LineageOT, and the ground-truth coupling. We visualized the remaining matrix elements in a UMAP embedding by connecting each RIM_parent cell to its confidently predicted descendants. To quantify method performance over the RIM_parent population, independent of the UMAP embedding and of any thresholding scheme, we computed the descendant error for RIM_parent cells, as described in our simulation study.

#### 2.3.2 Combining moslin with CellRank for fate mapping analysis

We focused on the ABpxp lineage (Strategy 1), and run moslin with the optimal hyperparameters identified in our grid search. We filtered out cells assigned a zero value in the marginal distributions to arrive at 6,476 cells used for this analysis. In the following, we used CellRank with default parameters if not indicated otherwise.

##### Transition matrix construction in CellRank

CellRank ^52^ is a fate mapping framework that was originally designed for RNA velocity ^66,67^ data. In version 2, it has been extended towards other data modalities, including time-series data. We make use of this extension here to construct a joint transition matrix *T* across all time-points for downstream CellRank analysis. Starting from an all-zero matrix *T*, containing cells from all time points, we execute the following steps:

1. First, we place moslin’s coupling matrices on the superdiagonal of *T* for transporting cells from early to late time points.
2. Second, we compute transition matrices within each time point based on gene expression similarity. We place these matrices on the diagonal of *T*.
3. Third, we compute a global transition matrix *T* ^*′*^ across all time points based on gene expression similarity. We combine *T* with *T* ^*′*^ with weights 0.9 and 0.1, respectively. This step improves matrix conditioning and yields the matrix *T* ^*′′*^.

We row-normalize *T* ^*′′*^ to arrive at the final CellRank transition matrix, which we interpret as a Markov chain. We simulated 200 random walks, each containing 500 steps, to visualize the predicted cell dynamics, starting from randomly selected 170 min cells.

##### Identifying terminal states and computing aggregated fate probabilities

We used CellRank’s GPCCA estimator ^68,69^ to compute 7 terminal states. We represented each terminal state by the 30 cells most confidently assigned to it. We aggregated individual terminal states to represent Ciliated neurons, Non-Ciliated neurons, and Glia and excretory cells, by combining the 30 cells identified per state. We computed absorption probabilities on the Markov chain towards these combined cell sets per terminal state group, and interpreted these as fate probabilities.

We used two-sided unequal variance Welch t-tests to asses whether fate probabilities were higher among pre-terminal cells for each terminal state group:

- for Ciliated neurons, we tested 647 pre-terminal ciliated neurons against 4,179 other preterminal and progenitor cells. We found *t* = 40.7, *P* = 6.4 · 10^−182^.
- for Non-ciliated neurons, we tested 890 pre-terminal non-ciliated neurons against 3,882 other pre-terminal and progenitor cells. We found *t* = 29.3, *P* = 3.0 · 10^−160^.
- for Glia and excretory cells, we tested 361 pre-terminal Glia and excretory cells against 4,227 other pre-terminal and progenitor cells. We found *t* = 82.2, *P* = 2.1 · 10^−255^.

##### Predicting driver genes

Using CellRank, we correlated each gene’s expression with the computed fate probabilities across all cells and subsetted to known *C. elegans* transcription factors ^70^ (TFs). We focused on the top 20 most strongly correlated TFs per terminal cell group and treated these as predicted driver TFs.

##### Computing a Palantir pseudotime

Using Palantir^71^, we computed a pseudotime from a randomly selected cell from the earliest embryo stage in our data. We used 30-nearest neighbors and sampled 1200 waypoint cells.

##### Visualizing expression trends in a heatmap

To visualize expression trends towards the non-ciliated neuron terminal state group, we selected the top 50 genes most strongly correlated with the corresponding fate probabilities (not subsetting to TFs). We imputed gene expression using MAGIC ^72^ and fitted Generalized Additive Models (GAMs) to each gene’s imputed expression as a function of the Palantir pseudotime, supplying non-ciliated neuron fate probabilities as cell-level weights to the loss function. Specifically, we used a spline basis and fitted GAMs with the mgcv package ^73^, through the CellRank interface.

### 2.4 Zebrafish heart regeneration (LINNAEUS)

The zebrafish heart regeneration dataset^74^ consists of hearts from 25 organisms; four uninjured hearts (ctrl), nine at three days after injury (3dpi), and seven at seven days after injury (7dpi). We use moslin to calculate couplings 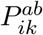, with *a* and *b* denoting datasets at consecutive timepoints. For ease of reading, we will suppress indices *a* and *b* in the following unless necessary.

#### 2.4.1 Mapping datasets

We embed the transcriptomic readout of all single cells with lineage information into a joint latent space using scVI ^45^, retaining the original cluster annotations. We calculate tree distances as shortest path distances along the original reconstructed trees. We use the moslin unbalanced FGW setting to calculate couplings between cells at consecutive timepoints. The standard parameters used are: *α* = 0.5, *ϵ* = 1*e* − 2, *τ*_*a*_ = 0.9, *τ*_*b*_ = 1, and *β* = 0.2. To understand the influence of the hyper-parameters on the performance we re-compute the mappings, changing a single parameter at a time within the following grids: *α* ∈ {0, 4, 0.6}, *ϵ* =∈ {5*e* − 3, 1*e* − 2, 1*e* − 1}, *τ*_*a*_ ∈ {0, 85, 0.95}, *τ*_*b*_ = 1, and *β* ∈ {0, 1, 0.3}.

In our calculations, we provide growth rates as initial marginals. To calculate growth rates, we use cell cycle marker genes typically used in single cell data ^75^ and the GSEA Hallmark apoptosis geneset (https://www.gsea-msigdb.org/gsea/msigdb/human/geneset/HALLMARK_APOPTOSIS.html). These are converted to their zebrafish orthologues using orthologues from Alliance, as previously described ^74^. Next, we use these two gene sets to calculate growth rates ^76^. For cells at 3dpi, that are in the regeneration process, we use the growth rates as calculated. However, cells at control are not in a regenerating heart and the calculated growth rates may not correlate with the actual injury response. Instead, we use cell type average growth rates as an approximation of the tendency of cell types to proliferate.

#### 2.4.2 Test for persistence of cell states

We expect that cells of the same, non-transient, type are persistent over time; cells of type *A* at time *t*_2_ should, for the most part, stem from cells of type *A* at time *t*_1_. This means moslin-computed couplings between those cells should be higher than those between cells of type *B* (with *B*≠*A*) at time *t*_1_ and cells of type *A* at time *t*_2_. To test this, we first select cell types at *t*_1_ and *t*_2_ with more than 10 cells that exist at both time points. We then define the distribution of couplings between cells of type *B* at *t*_1_ and cells of type *A* at *t*_2_ as

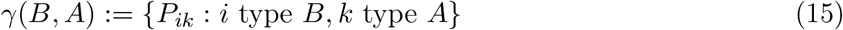

and perform a Welch’s t-test to calculate the significance level of the hypothesis

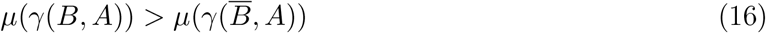

where 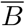 is the complement of *B*, i.e. all cells that are not type *B* (Fig. 5b), and *μ*(*γ*) is the mean of the population *γ*. Note that due to our requirement that every cell type at every time point contain at least 10 cells, all distributions here will have 100 or more datapoints, ample to assume the sample means are close to normal by the central limit theorem and therefore satisfy the normality assumption underlying a Welch’s t-test. The expectation of persistent non-transient cell types means that a significant test result for ⟨ *γ* (A,A) ⟩ > ⟨ *γ* (Ā,A) ⟩ is a true positive, and a significant test result for 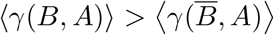 with *B*≠*A* is a false positive. With this formulation, we can create receiver operating characteristic (ROC) curves by iterating over the p-values from the t-tests and calculate the area under the ROC curve (AUC) value for a single combination of *t*_1_ and *t*_2_ datasets.

To create ROC curves for all control-3dpi and 3dpi-7dpi couplings, we perform this test between all *t*_1_ cell types and all *t*_2_ cell types within all combinations of datasets. We calculate AUCs to be 0.992 for control-3dpi and 0.984 for 3dpi-7dpi at hyper-parameter values *α* = 0.5, *β* = 0.2, *ϵ* = 0.01, *τ*_*a*_ = 0.9 (Fig. 5c and Subsection 1.3). To understand the influence of individual dataset couplings, we use the same procedure to calculate AUCs for all possible subsets of dataset combinations and plot the histogram of these AUCs (Fig. 5c, inset). Finally, to understand the influence of hyper-parameters *α, β, ϵ*, and *τ*_*a*_, we used the same procedure to calculate AUCs for couplings with different hyper-parameter values (Supp. Fig. 1). We observe no noticeable differences in AUCs.

Since the above described test follows a one-versus-one strategy, its AUC values may be inflated by a high amount of true negatives. We therefore implemented a variation of the cell type persistency test, following a one-versus-rest strategy. Here, we only test whether

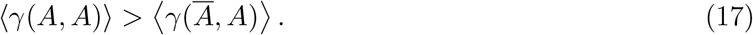

We calculate ROC curves as above and find areas under the curve of 0.9999 and 1 (up to eight decimals). We conclude that both in one-versus-one and one-versus-rest strategies, the cell type persistency test shows a very high performance of moslin.

#### 2.4.3 Calculating cellular flows

Given a coupling *P*_*ik*_ between a *t*_1_ dataset *a* and a *t*_2_ dataset *b*, cell type transitions from type *A* to type *B* can be quantified as

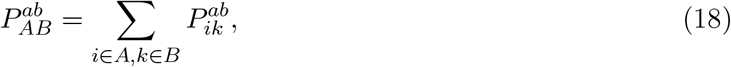

which satisfies Σ_*AB*_ *P*_*AB*_ = 1 since Σ_*ik*_ *P*_*ik*_ = 1. We construct weighted averages of these cell type transitions over all dataset combinations, weighing by the product of #*a* and #*b*, with #*a* the number of cells in *a*:

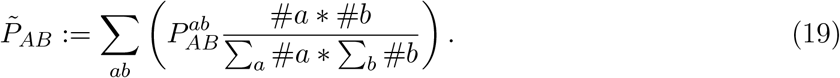

This definition satisfies 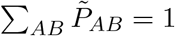.

We similarly obtain cell type frequencies at every timepoint by a weighted average of cell type frequencies 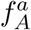 in each dataset *a* with weights #*a*:

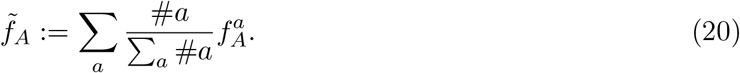

Again, 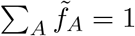 since 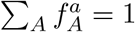 for each *a*.

To calculate the proportion *s*_*AB*_ of cells of type *A* becoming cells of type *B*, we divide 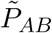 by the total mass outgoing from *A*:

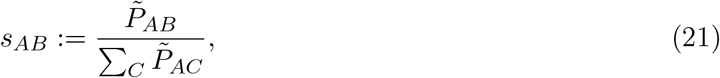

while the proportion *t*_*AB*_ of cells type *B* being generated by cells of type *A* is similarly calculated as

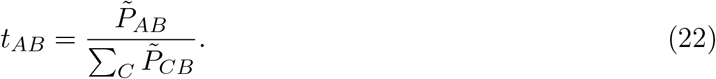

Finally, we subsampled the datasets used to calculate the proportions *s*_*AB*_, and then used the range of obtained values to determine confidence intervals. To reduce the amount of data roughly by half, we randomly selected three out of four control datasets, six out of nine 3dpi datasets and five out of seven 7dpi datasets, meaning 18 instead of 36 couplings between control and 3dpi datasets, and 30 instead of 63 couplings between 3dpi and 7dpi datasets. This method of random selection allows for a total of 7056 combinations:

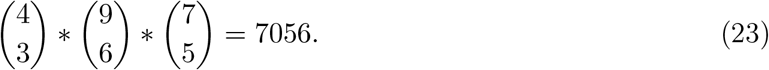

We explicitly calculated *s*_*AB*_ for all 7056 combinations to determine confidence intervals.

