## Supplementary Tables for "Mapping lineage-traced cells across time points with moslin"

**Supplementary Table 1: previously known driver genes in C. elegans**

Red, blue and green indicate genes moslin predicts to be driving ciliated, non-ciliated, and glia and excretory cells, respectively.

| Short name | WormBase ID | TF* | TF Family* | Involved in which cell type? | Reference |
|---|---|---|---|---|---|
| daf-19 | WBGene00000914 | Yes | RFX | Ciliated neurons | PMID: 10882127 |
| ceh-36 | WBGene00000457 | Yes | Homeobox | ASE, ASI, AWA and AWC sensory and ciliated neurons | PMID: 21041366, PMID: 15095973, PMID: 20150279, PMID: 24243022 , PMID: 14536063, PMID: 12952888, PMID: 32814896 |
| fkh-2 | WBGene00001434 | Yes | Fork_head | Ciliated neurons (AWB) | PMID: 17510633 |
| sptf-1 | WBGene00009743 | Yes | zf-C2H2 | Specific ciliated neurons (ASJ) | PMID: 25769980 |
| tub-1 | WBGene00006655 | Yes | Tub | Ciliated neurons | PMID: 31259686, PMID: 24646679 |
| daf-3 | WBGene00000899 | Yes | MH1 | Ciliated neurons (ADL) | PMID: 27351255 |
| fax-1 | WBGene00001400 | Yes | RXR-like | Interneurons (AVA, AVE) | PMID: 16183052 |
| zag-1 | WBGene00006970 | Yes | zf-C2H2 | Neurons, motor neurons | PMID: 12835394, PMID: 25474681, PMID: 12835395 |
| cnd-1 | WBGene00000561 | Yes | bHLH | Specific non-ciliated neurons, motor neurons | PMID: 32601060, PMID: 10976055 |
| ceh-24 | WBGene00000447 | Yes | Homeobox | Specific non-ciliated neurons (SIA and SIB) | PMID: 28244369 |
| unc-3 | WBGene00006743 | Yes | COE | Specific non-ciliated neurons, motor neurons | PMID: 25790851, PMID: 18817768 |
| ctbp-1 | WBGene00006424 | Yes | THAP | Motorneurons, Interneurons | PMID: 26480814, PMID: 35119366 |
| sdn-1 | WBGene00004749 | No | | Neuronal cell migration, Axon guidance | PMID: 16677626, PMID: 16176946 |
| syg-1 | WBGene00006365 | No | | Neurite growth in (non-ciliated) DD-type GABAergic neurons, Synapse formation for HSNL motor neurons | PMID: 35076532, PMID: 12628183, PMID: 15035988 |
| madd-4 | WBGene00009958 | No | | Synapse formation, axon guidance | PMID: 26028574, PMID: 32928959, PMID: 22014523, PMID: 25122090 |
| mig-13 | WBGene00003245 | No | | Neuron and neuroblast migration | PMID: 23784779, PMID: 26022293, PMID: 27780040 |
| flp-1 | WBGene00001444 | No | | Sensory and motor neurons, Non-ciliated AVK neurons, Interneuron signalling | PMID: 29293515, PMID: 16183052 |
| alr-1 | WBGene00044330 | Yes | Homeobox | Glial cells | PMID: 16055504 |
| pros-1 | WBGene00000448 | Yes | HPD | Glial cells, excretory cells | PMID: 27068465, PMID: 27402188, PMID: 23334499 |
| unc-130 | WBGene00006853 | Yes | Fork_head | Glial cells | PMID: 34423346 |
| ttx-1 | WBGene00006652 | Yes | Homeobox | Glial cells (AMsh glia) | PMID: 21350017 |
| ces-2 | WBGene00000469 | Yes | TF-bZIP | Excretory duct cell | PMID: 16310763, PMID: 12231624 |
| ref-2 | WBGene00004335 | Yes | zf-C2H2 | Interneurons (AIY) and excretory cells | PMID: 29442317, PMID: 19386265 |

| |
|---|
| * according to AnimalTFDB v4.0, see http://bioinfo.life.hust.edu.cn/AnimalTFDB4/#/ |
