## Supplementary Notes for "Mapping lineage-traced cells across time points with moslin"

April 6, 2023

### Moslin Supplementary Notes

### 1 Optimal transport (OT) to map cells across time-points

Following the success of previous methods[1] that link cells across time, moslin is based on optimal transport (OT) to reconstruct links between cross-sectional measurements. In this section, we give an introduction to OT[2] and highlight extensions that we use in moslin to incorporate lineage tracing data.

OT is an area of mathematics concerned with mapping distributions from one space to another; in our case, these distributions can be thought of as ensembles of sampled cells at different time points. The original notion of OT was introduced by the French mathematician Gaspard Monge in 1781 who considered the problem of moving a pile of sand to a hole where the shape of the pile and the hole are prescribed and a certain cost is associated with moving each grain of sand from source to target locations. This problem is known as the *Monge problem*; it leads to a constrained non-convex optimization problem that is not guaranteed to have a feasible solution in general[2]. Kantorovich[3] in 1942 famously proposed to relax the transport problem by allowing probability mass from a source destination to be split across several target destinations, thus evolving from deterministic *transport maps* to probabilistic *couplings*.

#### 1.1 The Kantorovich relaxation of OT

For probability vectors (equivalently, *histograms*) $\boldsymbol{a} \in \Delta_N$ and $\boldsymbol{b} \in \Delta_M$, define the corresponding discrete measures $\alpha$ and $\beta$ as

$$\alpha(\boldsymbol{x}) = \sum_{i=1}^{N} a_i \delta_{\boldsymbol{x}_i} \tag{1}$$

$$\beta(\boldsymbol{y}) = \sum_{j=1}^{M} b_i \delta_{\boldsymbol{y}_i}\ , \tag{2}$$

with respect to source and target locations, $(\boldsymbol{x}_i, \boldsymbol{y}_j) \in \mathcal{X} \times \mathcal{Y}$ for all $(i, j) \in \{1, ..., N\} \times \{1, ..., M\}$ where $\mathcal{X}, \mathcal{Y} \subseteq \mathbb{R}^D$. In our application, these represent $N$ sampled cells at $t_1$ and $M$ sampled cells at $t_1$.

OT over discrete probability measures seeks to find a *coupling matrix* $P \in \mathbb{R}_+^{N \times M}$, transporting mass from $\alpha$ to $\beta$ in a way that is optimal with respect to a cost function $c : \mathcal{X} \times \mathcal{Y} \to \mathbb{R}_+$ where $c(\boldsymbol{x}_i, \boldsymbol{y}_j)$ is the cost associated with moving a piece of unit mass from location $\boldsymbol{x}_i$ to $\boldsymbol{y}_j$. In our case, $\boldsymbol{x}_i$ and $\boldsymbol{y}_j$ can be thought of as cellular profiles in some embedded phenotypic space, e.g. the $D-$dimensional scVI[4] or PCA latent space, and $c$ quantifies biologically meaningful distance in that space.

The set of feasible couplings, given by those matrices $P$ that satisfy the marginal constraints imposed though the probability vectors $\boldsymbol{a}$ and $\boldsymbol{b}$, may be written as

$$U(\boldsymbol{a}, \boldsymbol{b}) := \left\{ P \in \mathbb{R}_+^{N \times M} : P\mathbf{1}_M = \boldsymbol{a},\ P^\top \mathbf{1}_N = \boldsymbol{b} \right\}\ , \tag{3}$$

for constant-one vectors $\mathbf{1}_N$ and $\mathbf{1}_M$ of lenghts $N$ and $M$, respectively. In our case, the marginals $\boldsymbol{a}$ and $\boldsymbol{b}$ describe distributions over sampled cell states at early and late time points, respectively. They may be chosen to be uniform; alternatively, they can reflect any cell-level prior such as cellular growth and death.

With these definitions at hand, we can state the Kantorovich relaxation of the OT problem as follows:

$$\mathcal{L}_c(\alpha, \beta) := \min_{P \in U(\boldsymbol{a}, \boldsymbol{b})} \langle C, P \rangle\ , \tag{4}$$

for cost matrix $C \in R_+^{N \times M}$ with $C_{ij} = c(\boldsymbol{x}_i, \boldsymbol{y}_j)$ and $\langle C, P \rangle := \sum_{ij} C_{ij} P_{ij}$. Following Peyré et al.[2], we make the dependency of the OT problem $\mathcal{L}_c$ on the cost function $c$ explicit. In case the cost function $c$ is related to a proper distance metric $d$ via $c(\boldsymbol{x}_i, \boldsymbol{y}_j) = d^p(\boldsymbol{x}_i, \boldsymbol{y}_j)$, i.e. the cost is given by the $p$-th power of a distance metric, we define the *$p$-Wasserstein distance* as

$$W_p(\alpha, \beta) = \mathcal{L}_{d^p}(\alpha, \beta)^{1/p}, \tag{5}$$

where $\mathcal{L}_{d^p}$ is defined as in Equation (4). We refer to Peyré et al. [2] for an overview and to Chen et al. [5] for a single-cell application of Wasserstein distances to describe patient-level variation in terms of single-cell gene expression.

**Practical considerations.** The objective function defined in Equation (4) is linear with constraints given by the $N + M$ equality constraints imposed though $U(\boldsymbol{a}, \boldsymbol{b})$, thus it defines a convex linear program whose solution is in general non-unique. Various strategies have been suggested to solve Equation (4), among them network flow solvers and the auction algorithm [6], however, all of these remain limited to finding a solution in time $\mathcal{O}(N^3)$ for $M = N$, omitting logarithmic factors. This poses a scalability issue for applications in current single-cell genomics datasets which frequently contain hundreds of thousands of cells. Further, practical limitations include the difficulty to adapt these algorithms to run on GPUs and to be differentiable.

#### 1.2 Entropic regularization and the Sinkhorn algorithm

To overcome these practical limitations, consider the following entropically regularized [7] variant of the OT problem,

$$\mathcal{L}_c^{\epsilon}(\alpha, \beta) \coloneqq \min_{P \in U(\boldsymbol{a}, \boldsymbol{b})} \langle P, C \rangle - \epsilon H(P), \tag{6}$$

for regularization strength $\epsilon > 0$ and entropy term

$$H(P) \coloneqq -\sum_{ij} P_{ij} \left(\log P_{ij} - 1\right). \tag{7}$$

In contrast to the unregularized problem, Equation (6) is $\epsilon$-strongly convex and thus possesses a unique global optimum. Further, for $\epsilon \to 0$ and $\epsilon \to \infty$, Peyré et al. [2] and Cominetti and Martín [8] show the following asymptotic results:

$$\mathcal{L}_c^{\epsilon}(\alpha, \beta) \to \mathcal{L}_c \text{ for } \epsilon \to 0, \tag{8}$$

$$P_{\epsilon} \to \boldsymbol{a}\boldsymbol{b}^{\top} \text{ for } \epsilon \to \infty, \tag{9}$$

where $P_{\epsilon}$ refers to the solution of the regularized problem with regularization parameter $\epsilon$. In particular, these results show that for small $\epsilon$, the regularized problem approximates the unregularized problem while for large $\epsilon$, the coupling converges to the outer product of the two marginals, which is closely related to the *Maximum Mean Discrepancy* distance commonly used in generative models for distribution matching, see Lotfollahi et al. [9] for a single-cell application. To solve the regularized OT problem of Equation (6), we reproduce the following proposition from Peyré et al. [2]:

**Proposition 1.1** (Solution to the regularized OT problem)**.** *The unique solution to the regularized OT problem introduced in Equation (6) can be written as*

$$P_{ij} = u_i K_{ij} v_j \quad \forall (i, j) \in \{1, ..., N\} \times \{1, ..., M\}, \tag{10}$$

*for the associated Gibbs kernel* $K_{ij} = \exp\left(-C_{ij}/\epsilon\right)$ *and scaling variables* $(\boldsymbol{u}, \boldsymbol{v}) \in \mathbb{R}_+^N \times \mathbb{R}_+^M$ *to be inferred.*

*Proof.* For a proof, see Peyré et al. [2]. □

**Sinkhorn algorithm.** The form of the solution outlined in Proposition 1.1 can be used to construct an algorithm that iterates between scaling the rows and the columns of a candidate matrix. Impose therefore the marginal constraints of the feasible set $U(\boldsymbol{a}, \boldsymbol{b})$ on a solution in the form of Proposition 1.1,

$$\operatorname{diag}(\boldsymbol{u})\, K \operatorname{diag}(\boldsymbol{v})\, \mathbf{1}_M = \boldsymbol{a}\,, \tag{11}$$

$$\operatorname{diag}(\boldsymbol{v})\, K^\top \operatorname{diag}(\boldsymbol{u})\, \mathbf{1}_N = \boldsymbol{b}\,, \tag{12}$$

which can be simplified to give

$$\boldsymbol{u} \odot (K\boldsymbol{v}) = \boldsymbol{a}\,, \tag{13}$$

$$\boldsymbol{v} \odot (K^\top \boldsymbol{u}) = \boldsymbol{b}\,, \tag{14}$$

with $\odot$ denoting elements-wise multiplication. Iteratively solving these equations gives rise to *Sinkhorn's algorithm*,

$$\boldsymbol{u}^{(l+1)} := \frac{\boldsymbol{a}}{K\boldsymbol{v}^{(l)}}\,, \tag{15}$$

$$\boldsymbol{v}^{(l+1)} := \frac{\boldsymbol{b}}{K^\top \boldsymbol{u}^{(l+1)}}\,, \tag{16}$$

where the division is applied element-wise. Yule[10] originally suggested iterations of this form, Sinkhorn[11] proofed their convergence and Cuturi[7] suggested applying the algorithm to solve entropically regularized OT problems which gives rise to a differentiable solution. Note that Sinkhorn's algorithm is well suited to run on GPUs since it only relies on matrix vector products.

Cuturi et al.[12] recently used JAX[13] to implement Sinkhorn's algorithm in their Optimal Transport Tools (OTT) software package; OTT thus allows for just-in-time compilation, GPU acceleration, online cost function evaluation, and automatic differentiation. Moslin is implemented in moscot, which connects OTT to the single-cell world. In contrast to the linear programming algorithms introduced above, Sinkhorn runs in time $\mathcal{O}(N^2)$ for $N = M$, omitting logarithmic factors. Thus, solving the regularized (Equation (6)) rather than the unregularized (Equation (4)) OT problem offers both practical and theoretical advantages including improved statistical convergence[2,14].

#### 1.3 OT extensions: unbalancedness and (Fused) Gromov-Wasserstein (FGW)

The moslin model requires two further extensions of the regularized OT problem, we follow Peyré et al.[2] in their presentation. First, cells proliferate and die while they differentiate which we include in the marginal distributions $\boldsymbol{a}$ and $\boldsymbol{b}$. However, as the rates of cellular growth and death are difficult to estimate based on scRNA-seq data, we relax the marginal constraints which leads to *unbalanced optimal transport*.

Second, lineage tracing information in ICE is only comparable within one time point as barcodes are generated independently in each replicate. Nevertheless, it is reasonable to assume that pairwise lineage similarity is conserved on average; across time points, cell pairs with similar lineage distances are more likely to belong together compared to cell pairs with very different lineage distances. Thus, we allow for pairwise comparisons of lineage information only within each time point which leads to *Gromov-Wasserstein OT* (GW).

**Unbalanced OT.** To relax the marginal constraints, consider a generalization of the regularized OT problem by adding divergences $D_\phi$ between marginal constraints and row/columns sums of $P$ to the objective function,

$$\mathcal{L}_c^{\epsilon,\tau}(\alpha, \beta) := \min_{P \in \mathbb{R}_+^{N \times M}} \langle C, P \rangle + \rho_a D_\phi(P\mathbf{1}_M | \boldsymbol{a}) + \rho_b D_\phi(P^\top \mathbf{1}_N | \boldsymbol{b}) - \epsilon H(P)\,, \tag{17}$$

where the parameters $\rho_a, \rho_b > 0$ control the weight given to the soft marginal constraints[2,15]. We follow OTT[12] and use the parametrization $\rho_{a,b} := \frac{\epsilon \tau_{a,b}}{1-\tau_{a,b}}$ for $\tau_{a,b} \in (0, 1)$.

This is a generalization of the original entropic OT problem of Equation (6) as in the limit $\tau_1, \tau_2 \to 1$, i.e., $\rho_a, \rho_b \to \infty$, one recovers the original problem. A generalized version of the Sinkhorn algorithm may be applied to solve Equation (17); of particular importance in practical applications[1] has been the case $D_\phi(\cdot|\cdot) = \mathrm{KL}\left[\cdot \,||\, \cdot\right]$ for which the Sinkhorn updates read

$$\boldsymbol{u}^{(l+1)} := \left(\frac{\boldsymbol{a}}{K\boldsymbol{v}^{(l)}}\right)^{\tau_a}, \tag{18}$$

$$\boldsymbol{v}^{(l+1)} := \left(\frac{\boldsymbol{b}}{K^\top \boldsymbol{u}^{(l+1)}}\right)^{\tau_b}. \tag{19}$$

We refer to Liero et al.[15] for a treatment of the theory behind unbalanced OT and to Chizat et al.[16] for the derivation of practical algorithms.

**Unbalancedness for approximate growth rates and cellular sampling.** The generalized Sinkhorn algorithm provides an efficient solution to unbalanced OT problems which arise in moslin when allowing for cellular growth and death at an uncertain rate $g$, or when considering the stochastic effects of sampling cells from a population (Methods).

**Gromov-Wasserstein optimal transport (GW).** In standard OT, we assume that point clouds $\{\boldsymbol{x}_1, ..., \boldsymbol{x}_N\}$ and $\{\boldsymbol{y}_1, ..., \boldsymbol{y}_M\}$ corresponding to bins in the histograms $\boldsymbol{a}$ and $\boldsymbol{b}$ may be compared using the cost function $c(\cdot, \cdot)$, i.e. $\mathcal{X}$ and $\mathcal{Y}$ correspond to the same metric space, giving rise to the cost matrix $C$. To relax this assumption, consider a situation where vectors $\{\boldsymbol{x}_i\}_{i=1}^N$ may be compared using the cost function $c^{\mathcal{X}}$ and vectors $\{\boldsymbol{y}_i\}_{i=1}^M$ may be compared using the cost function $c^{\mathcal{Y}}$, but no direct comparisons of vectors in $\mathcal{X}$ and $\mathcal{Y}$ are possible. Using these cost functions, we define the entropically regularized GW problem,

$$\mathcal{L}^{\epsilon}_{L,c^{\mathcal{X}},c^{\mathcal{Y}}}(\alpha, \beta) := \min_{P \in U(\boldsymbol{a},\boldsymbol{b})} \sum_{ijkl} L\left(C^{\mathcal{X}}_{ij}, C^{\mathcal{Y}}_{kl}\right) P_{ik} P_{jl} - \epsilon H(P), \tag{20}$$

for cost matrices $C^{\mathcal{X}} \in R_+^{N \times N}$, $C^{\mathcal{Y}} \in R_+^{M \times M}$ with $C^{\mathcal{X}}_{ij} = c^{\mathcal{X}}(\boldsymbol{x}_i, \boldsymbol{x}_j)$, $C^{\mathcal{Y}}_{kl} = c^{\mathcal{Y}}(\boldsymbol{y}_k, \boldsymbol{y}_l)$ and distance metric $L$. The solution to (the unregularized version of) Equation (20) defines the *Gromov-Wasserstein distance* between two metric spaces, each equipped with a probability distribution. This distance has been introduced by Mémoli[17] as an extension to the Gromov-Hausdorff distance[18], combined with entropic regularization by Peyré et al.[19], Solomon et al.[20] and used in the single-cell field e.g. for data integration across modalities[21]. Equation (20) defines a non-convex, constrained, smooth optimization problem in $P$; before discussing its optimization, we introduce one further generalization.

**Fused Gromov-Wasserstein optimal transport.** In the moslin model, while lineage information is only comparable within one time point, molecular similarity is comparable across time points. Thus, we encounter a problem that possesses characteristics of both OT (Equation (6)) and GW (Equation (20)). Problems of this kind require a combined objective function where some sampled features may be compared across spaces while others may only be compared within one space. This kind of problem is known as *entropically-regularized Fused Gromov-Wasserstein* (FGW), which has previously been employed in the single-cell field to reconstruct spatial cell arrangement[22]. The problem is defined via

$$\mathcal{L}^{\epsilon,\alpha}_{L,c^{\mathcal{X}},c^{\mathcal{Y}},c}(\alpha, \beta) := \min_{P \in U(\boldsymbol{a},\boldsymbol{b})} \alpha \sum_{ijkl} L\left(C^{\mathcal{X}}_{ij}, C^{\mathcal{Y}}_{kl}\right) P_{ik} P_{jl} + (1-\alpha) \sum_{ik} C_{ik} P_{ik} - \epsilon H(P), \tag{21}$$

where $\alpha \in [0,1]$ controls the weight given to the OT versus GW terms, the within-space cost functions $c^{\mathcal{X}}$ and $c^{\mathcal{Y}}$ are defined for these features $\{\boldsymbol{x}_i\}_{i=1}^N$ and $\{\boldsymbol{y}_i\}_{i=1}^M$ which may not be compared across spaces and the across-space cost function $c$ is defined for these features $(\boldsymbol{x}'_i, \boldsymbol{y}'_i)$ which may be compared across spaces[23]. For $\alpha = 1$, we recover the GW problem introduced above.

Introducing the 4-tensor[19],

$$\mathcal{T}(C^{\mathcal{X}}, C^{\mathcal{Y}})_{ijkl} := L\left(C^{\mathcal{X}}_{ik}, C^{\mathcal{Y}}_{jl}\right), \tag{22}$$

allows us to rewrite Equation (21) in shorter form,

$$\mathcal{L}^{\epsilon,\alpha}_{L,c^{\mathcal{X}},c^{\mathcal{Y}},c}(\alpha,\beta) := \min_{P \in U(\boldsymbol{a},\boldsymbol{b})} \alpha \left\langle \mathcal{T}(C^{\mathcal{X}}, C^{\mathcal{Y}}) \otimes P, P \right\rangle + (1-\alpha)\left\langle C, P \right\rangle - \epsilon H(P) \tag{23}$$

with tensor multiplication defined via

$$(\mathcal{T} \otimes P)_{ij} := \sum_{kl} \mathcal{T}_{ijkl} P_{kl}\,. \tag{24}$$

This tensor product may be computed in time $\mathcal{O}(N^3)$ for $M = N$ for a class of separable loss functions $L$, including $l_2$ loss and the KL divergence[19].

**FGW optimization.** Following Peyré et al.[19], we use projected gradient descent with iterations

$$P^{(l+1)} = \mathrm{Proj}^{\mathrm{KL}}_{U(\boldsymbol{a},\boldsymbol{b})}\left(P^{(l)} \odot e^{-\tau \nabla J(P)|_{P^{(l)}}}\right), \tag{25}$$

where $\mathrm{Proj}^{\mathrm{KL}}_{U(\boldsymbol{a},\boldsymbol{b})}(\tilde{P}) = \mathrm{argmin}_{P \in U(\boldsymbol{a},\boldsymbol{b})} \sum_{ij} P_{ij} \log\left(P_{ij}/\tilde{P}_{ij}\right)$ is a KL projection operator, $\tau$ is a step size, $J$ is the FGW objective function defined in Equation (21) and $\odot$ denotes element-wise multiplication. The gradient of the objective function may be written as

$$\nabla J = (1-\alpha)C + \alpha \mathcal{T}\left(C^{\mathcal{X}}, C^{\mathcal{Y}}\right) \otimes P\,, \tag{26}$$

while the KL projection can be solved via an OT problem[24],

$$\mathrm{Proj}^{\mathrm{KL}}_{U(\boldsymbol{a},\boldsymbol{b})}(\tilde{P}) = \operatorname*{argmin}_{P \in U(\boldsymbol{a},\boldsymbol{b})} \left\langle -\epsilon \log \tilde{P}, P \right\rangle - \epsilon H(P)\,. \tag{27}$$

Using Equation (26), Equation (27) and setting $\tau = 1/\epsilon$, we can re-write the update rule of Equation (25) as

$$P^{(l+1)} = \operatorname*{argmin}_{P \in U(\boldsymbol{a},\boldsymbol{b})} \left\langle (1-\alpha)C + \alpha \mathcal{T}\left(C^{\mathcal{X}}, C^{\mathcal{Y}}\right) \otimes P^{(l)}, P \right\rangle - \epsilon H(P)\,, \tag{28}$$

which is the entropically regularized OT problem of Equation (6), solved efficiently at each iteration for an evolving cost matrix using the Sinkhorn algorithm[7,19,22]. The algorithm outlined here is applicable to both GW ($\alpha = 1$) and FGW ($\alpha \in (0,1)$) settings. The major computational bottleneck is the update of the tensor product of Equation (24) required at each update of Equation (28), which runs in time $\mathcal{O}(N^3)$ for $N = M$.

**Unbalanced FGW optimal transport.** We showed in Equation (17) how OT may be relaxed to allow deviations from the exact marginals $\boldsymbol{a}, \boldsymbol{b}$ to accommodate uncertain cellular growth and death-rates; we may use a similar trick for FGW by replacing the hard marginal constraints with soft quadratic $\phi$ divergences $D^{\otimes}_{\phi}$. This has been explored by Séjourné et al.[25] where the authors introduce the *Unbalanced Gromov-Wasserstein* (UGW) divergence and show that it upper-bounds a proper distance metric between metric spaces equipped with a positive measure.

Combining unbalancedness with FGW leads to our final optimization problem which captures the various problem aspects:

- a Wasserstein term to compare cells across time points using e.g. molecular similarity[1].
- a Gromov-Wasserstein[17,19] term to compare cells within time points using e.g. lineage information in ICE.
- unbalancedness[16,26] to handle stochastic cell sampling, and uncertain growth- and death-rates.
- entropic regularization for improved computational[2,7] and statistical properties[14].

Thus, moslin's OT formulation allows the method to flexibly capture the various problem aspects of linking lineage-traced cells across time points.

[7] Marco Cuturi. Sinkhorn distances: Lightspeed computation of optimal transport. *Advances in neural information processing systems*, 26, 2013.
