## Supplementary Figures for "Mapping lineage-traced cells across time points with moslin"

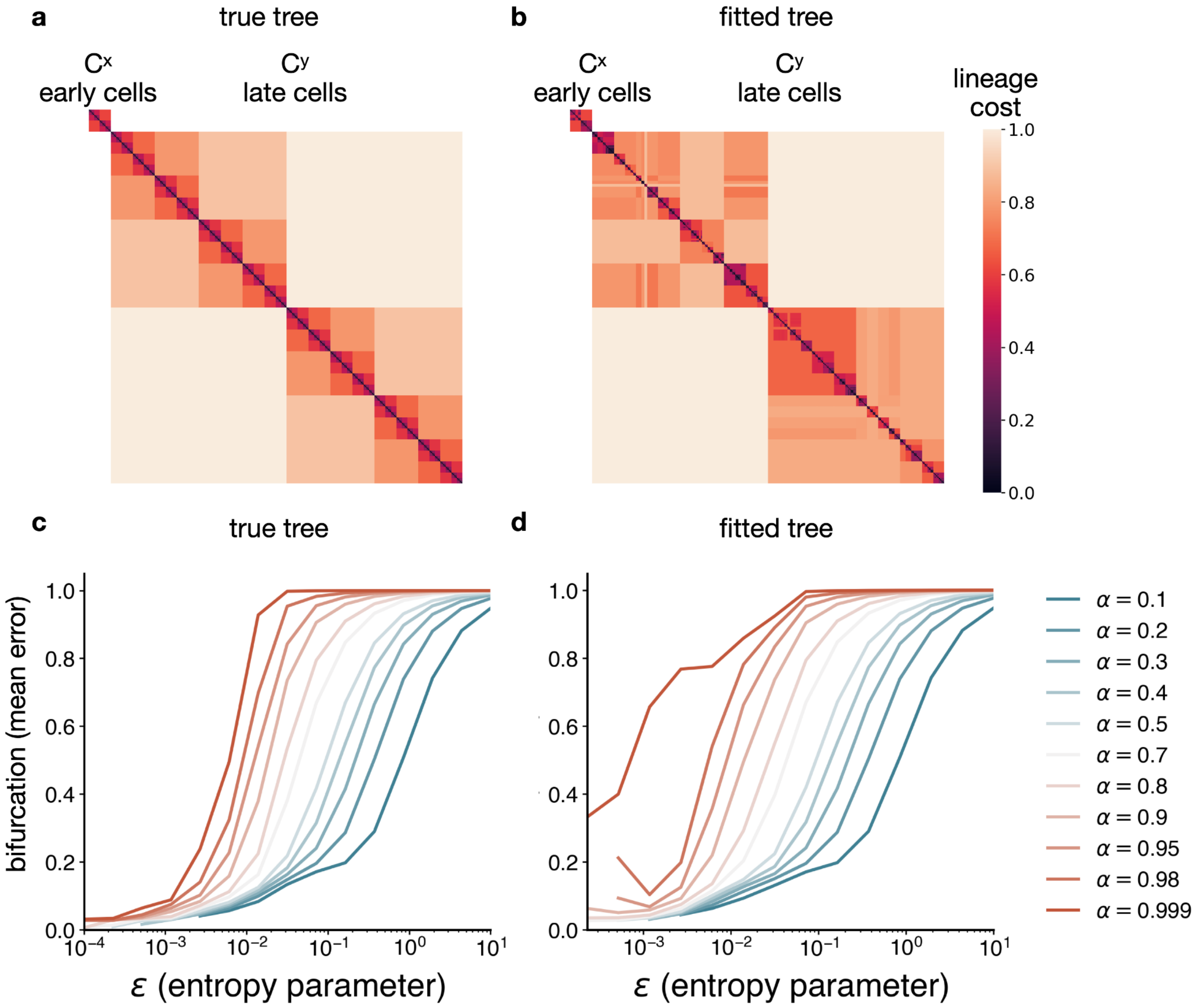


**Suppl. Fig. 1 | The bifurcation trajectory as an illustrative example of moslin's performance.**

**a,b.** Heatmaps, visualizing the lineage distance cost matrices $C^X$ and $C^Y$, corresponding to lineage distance between early and late cells, respectively, as used by moslin in the bifurcation trajectory setting for the true tree (**a**) and the fitted tree (**b**). Higher cost values imply cells are further apart in the tree. **c,d.** Line plots, illustrating moslin's bifurcation-trajectory grid search results for the true tree (**c**) and the fitted tree (**d**), across a range of $\alpha$ values. The x- and y-axis display the entropy parameter $\epsilon$ and the mean over ancestor and descendant errors, respectively.

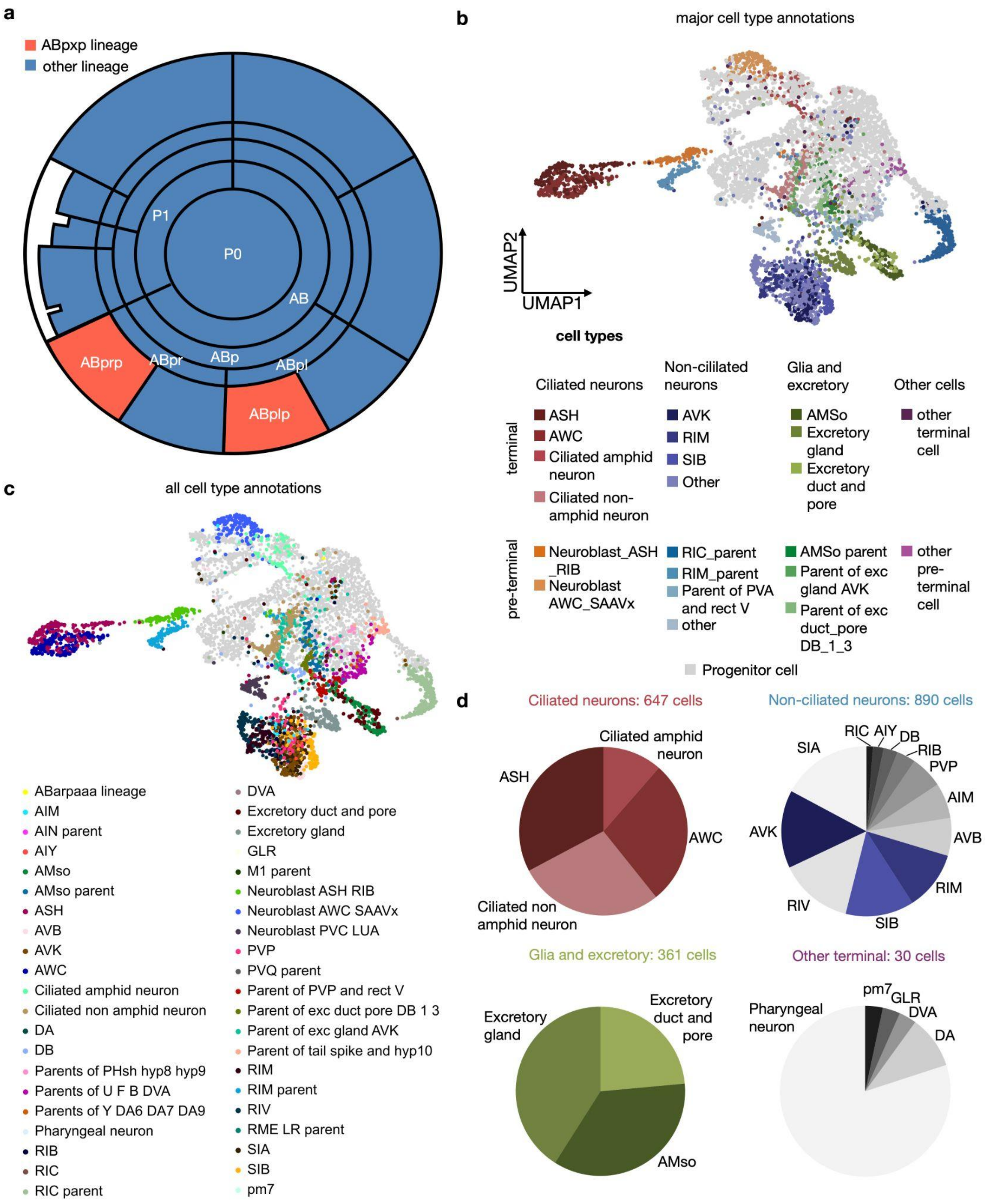


**Suppl. Fig. 2 | Cluster and lineage labels for the *C. elegans* data.**

**a.** Conceptual illustration of the *C. elegans* ABpxp lineage, adjusted from ref.[7]. **b,c.** UMAPs for cells of the ABpxp lineage, colored by cluster annotations for Ciliated neurons, Non-ciliated neurons and Glia and excretory cells (**b**), and all cluster annotations from ref.[7] (**c**). **d.** Pie charts visualizing the makeup of terminal Ciliated neurons, Non-ciliated neurons, Glia and excretory, and other terminal cells.

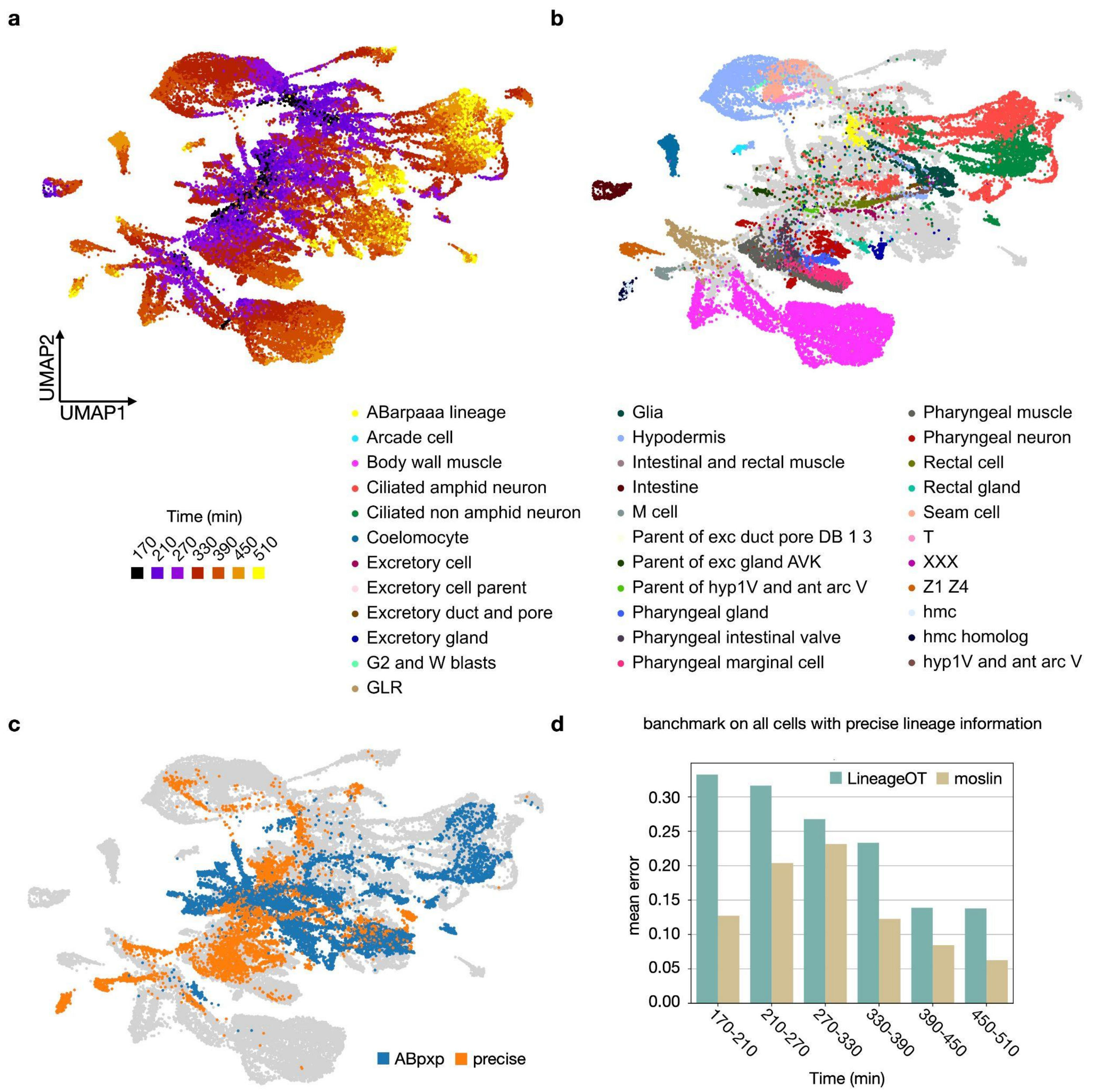


**Suppl. Fig. 3 | Benchmark on all cells with precise lineage information.**

**a-c.** UMAPs of 45,732 *C.elegans* cells with at least partial lineage information, colored by time point (**a**), cell type (**b**), and the subsets used in this work for benchmarking (**c**). **d.** Bar chart of the mean error for moslin and LineageOT[21] across time points as in Fig. 3b, but for all cells with precise lineage information shown in (**c**).

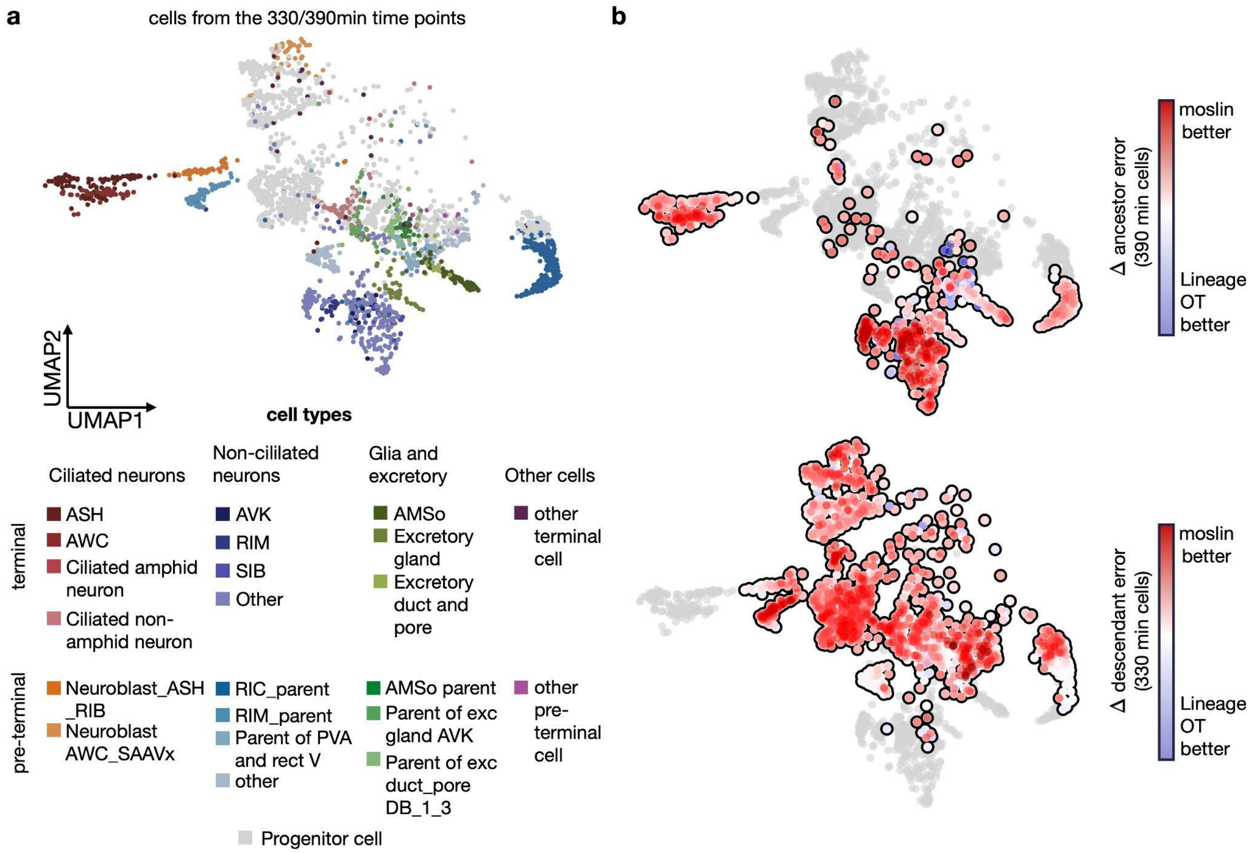


**Suppl. Fig. 4 | Ancestor and descendant error on the 330/390min time point pair.**

**a,b.** UMAPs of 330/390min cells of the ABpxp lineage, colored by cluster annotations (**a**) and the difference in ancestor and descendant error between moslin and LineageOT (**b**).

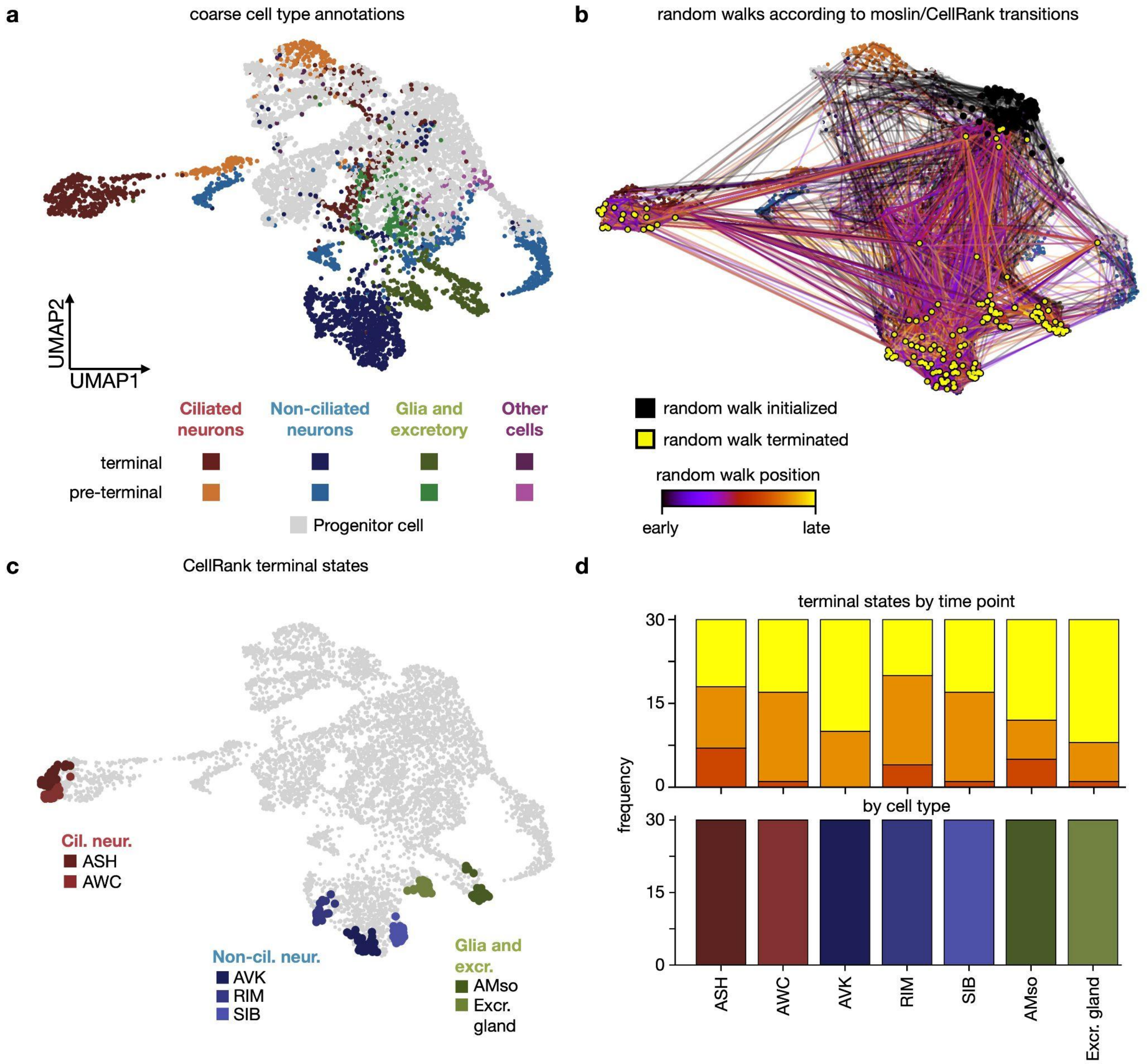


**Suppl. Fig. 5 | Random walks and terminal states.**

**a,b.** UMAP embeddings of the ABpxp lineage, colored by coarse cell-type annotations. **b.** Additionally, we show random walks on the moslin/CellRank computed transition matrix, initialized in 170 min cells (Methods). Random walks progress towards the expected terminal populations. **c.** UMAP, showing the top 30 cells per moslin/CellRank[22] computed terminal state. **d.** Bar charts, showing how the top 30 cells per terminal state distribute across time points and clusters.

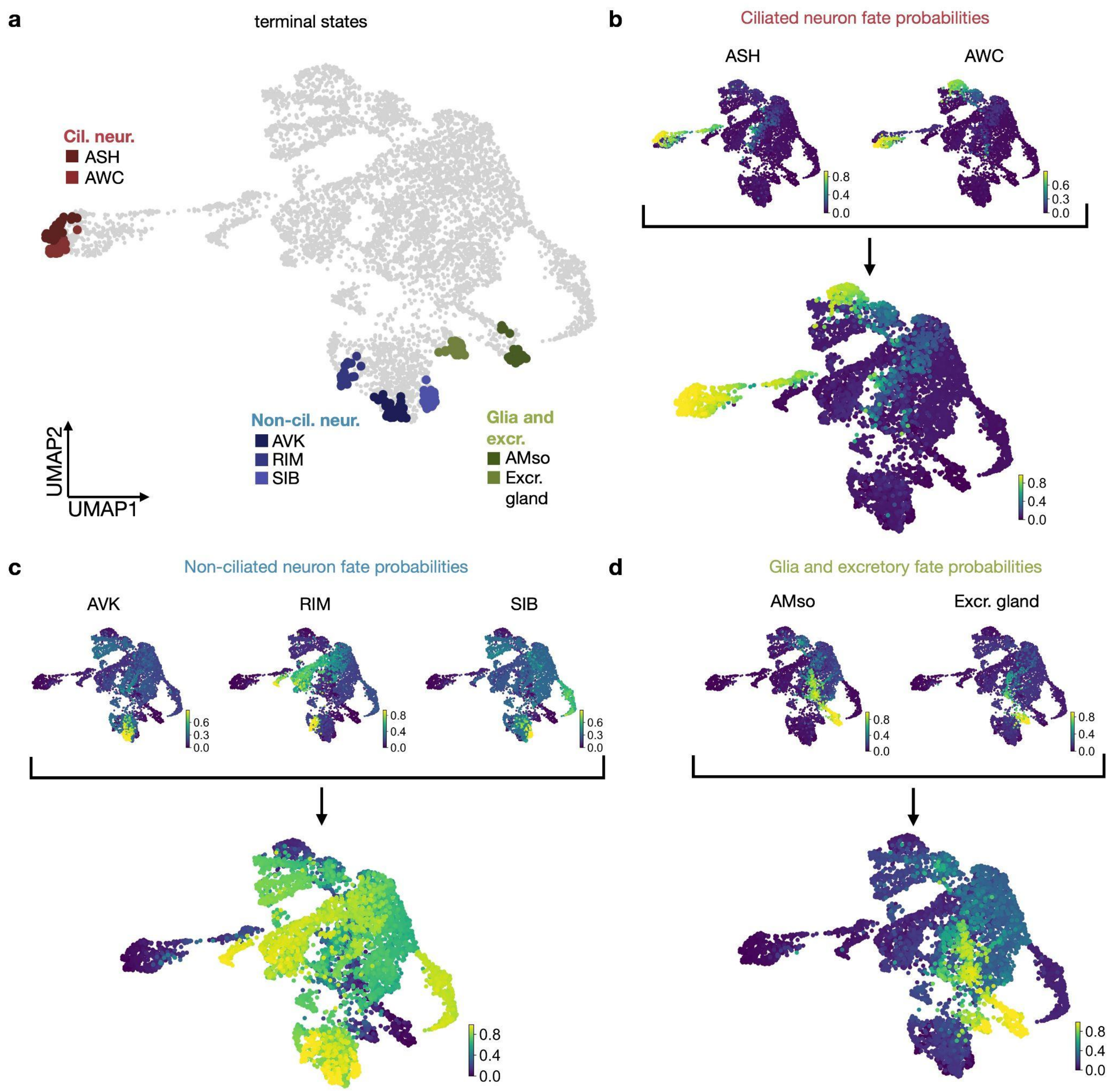


**Suppl. Fig. 6 | Aggregating fate probabilities towards the main terminal groups.**

**a.** UMAP of the ABpxp lineage, with CellRank-computed terminal states indicated, as in Fig. 3. **b-d.** Aggregation of individual fate probabilities towards Ciliated neurons (**b**), Non-ciliated neurons (**c**), and Glia and excretory cells (**d**).

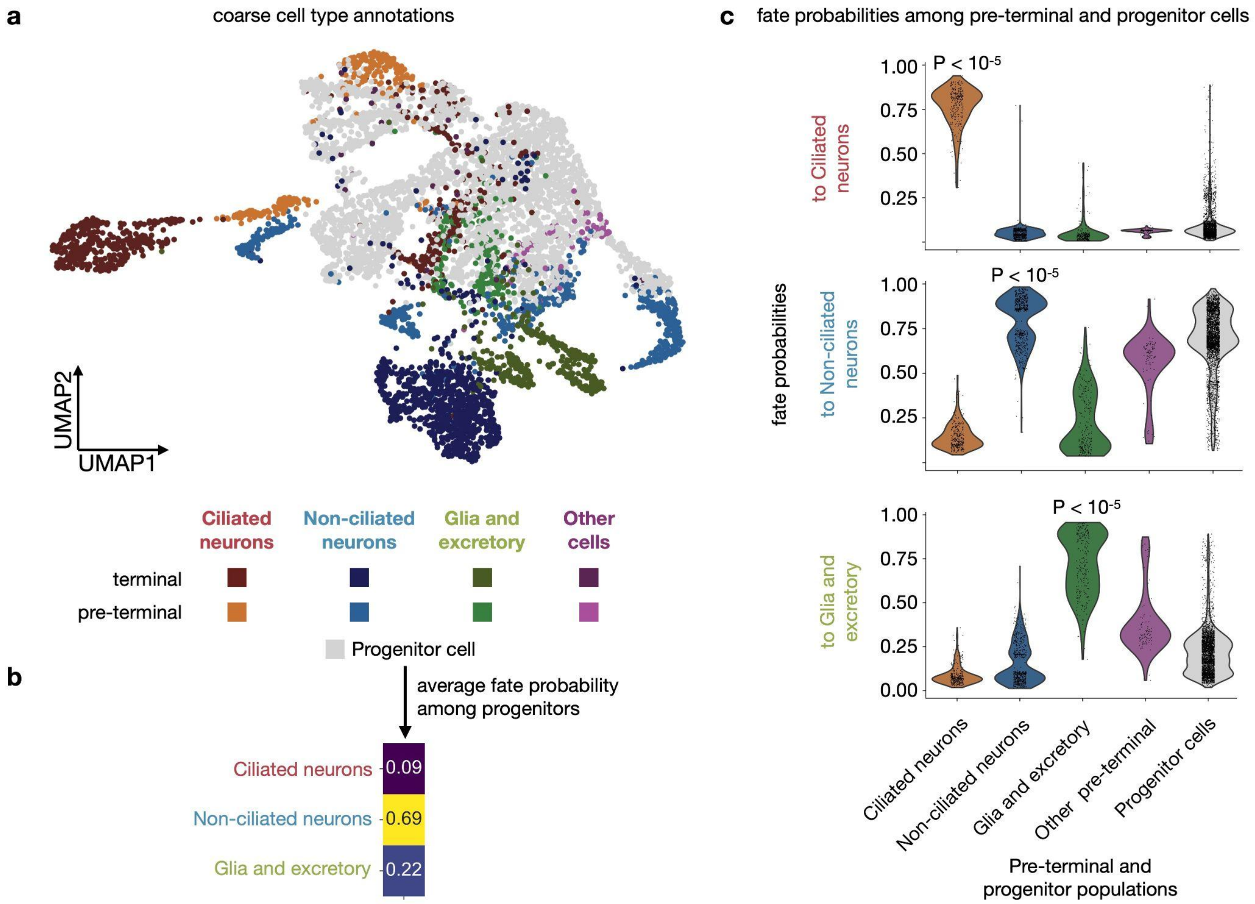


**Suppl. Fig. 7 | Fate probabilities are higher in corresponding pre-terminal populations.**

**a.** UMAP of the ABpxp lineage, colored by coarse cluster annotations. **b.** Heatmap, displaying average fate probability among progenitors cells. **c.** Violin plots over the distribution of fate probabilities towards Ciliated neurons (top), Non-ciliated neurons (center), and Glia and excretory cells (bottom), grouped by pre-terminal cell states as shown in (**a**). For each terminal state, fate probabilities in the corresponding pre-terminal group are significantly higher compared to all other groups (two-sides Welch's t-tests; Methods)

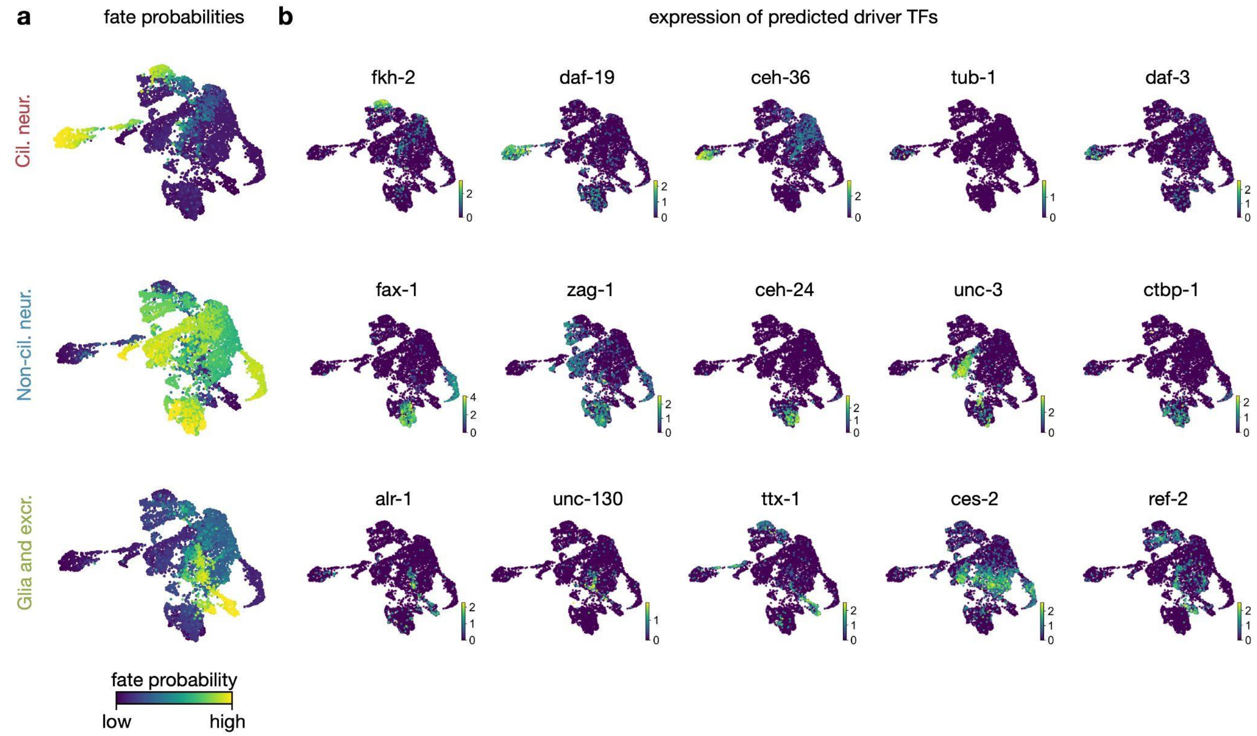


**Suppl. Fig. 8 | Predicted driver TFs for Ciliated and Non-ciliated neurons, Glia and excretory cells.**

**a,b.** UMAPs of the ABpxp lineage, colored by moslin/Cellrank-computed fate probabilities (**a**) and the expression of predicted driver TFs (**b**). Each TF shown here is among the 20 most correlated TFs with the corresponding fate probabilities and has previously been reported to be important for the corresponding developmental trajectory (Methods and Supplementary Table 1).

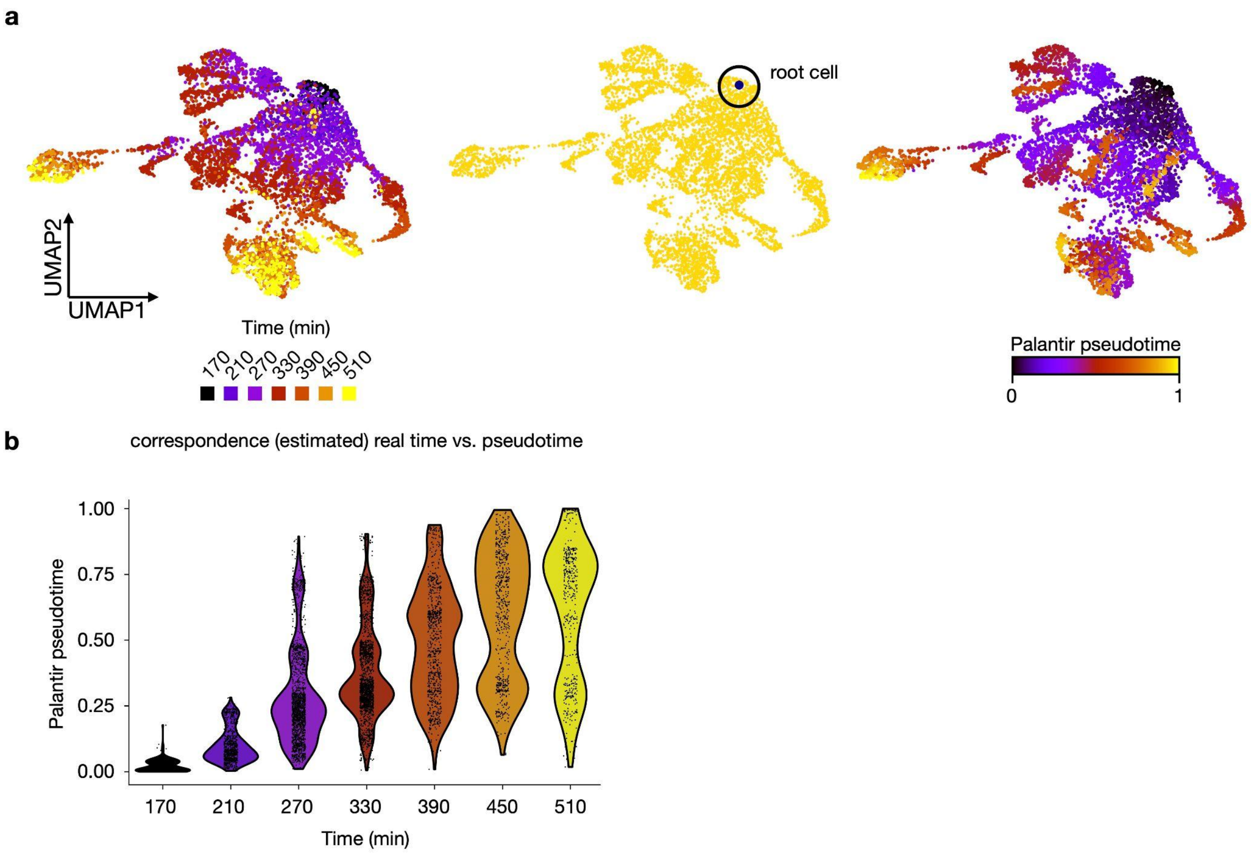


**Suppl. Fig. 9 | Computing a pseudotime using Palantir.**

**a.** UMAPs of the ABpxp lineage, colored by (from left): estimated real time points (Methods), a 170min cell passed to Palantir as root cell, and the Palantir pseudotime[53]. **b.** Violin plot over the Palantir pseudotime, grouped by estimated real time points.

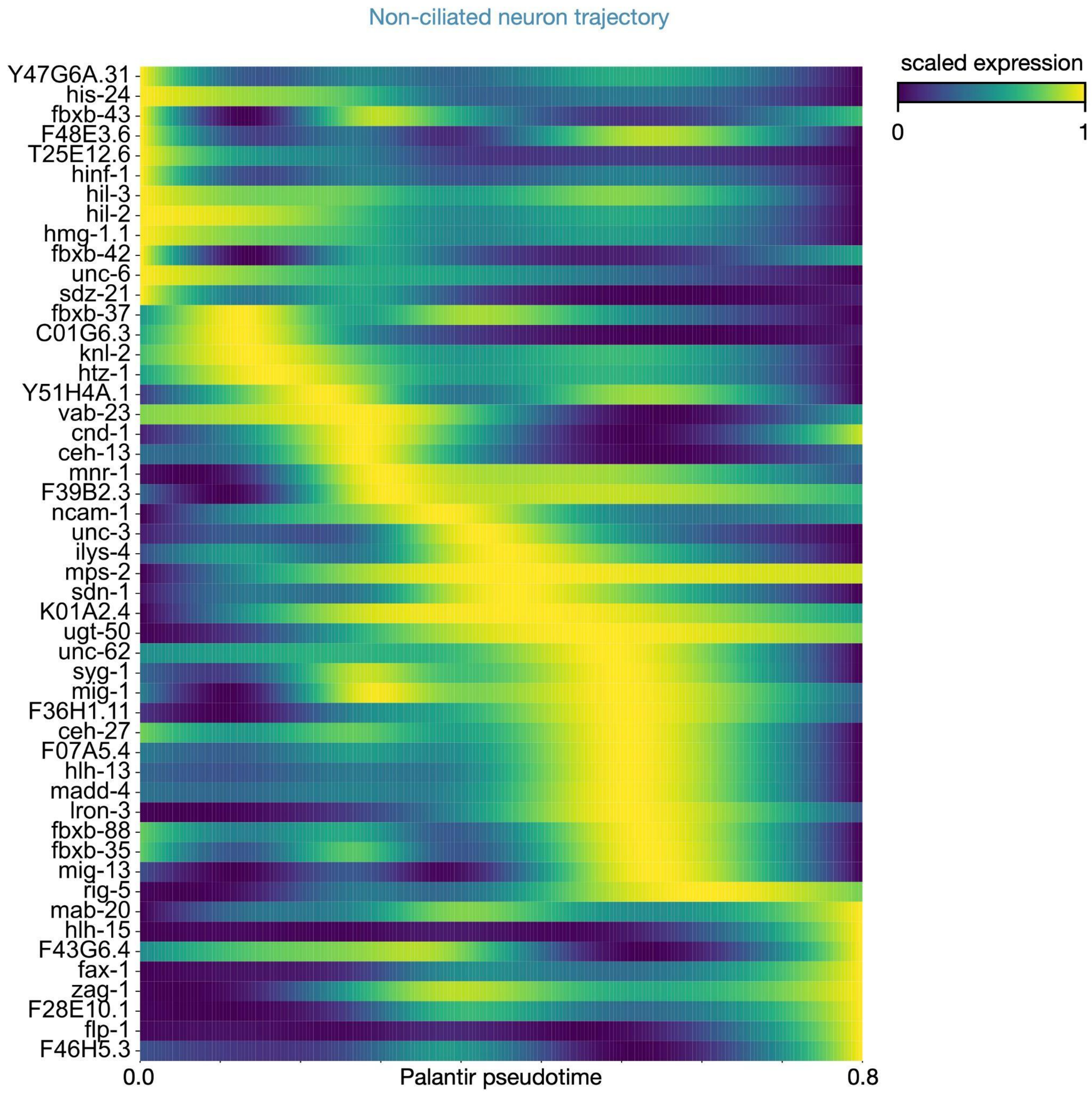


**Suppl. Fig. 10 | Complete heatmap of smoothed gene expression along the non-ciliated neuron trajectory.**

As in Fig. 3f, with all gene names included.

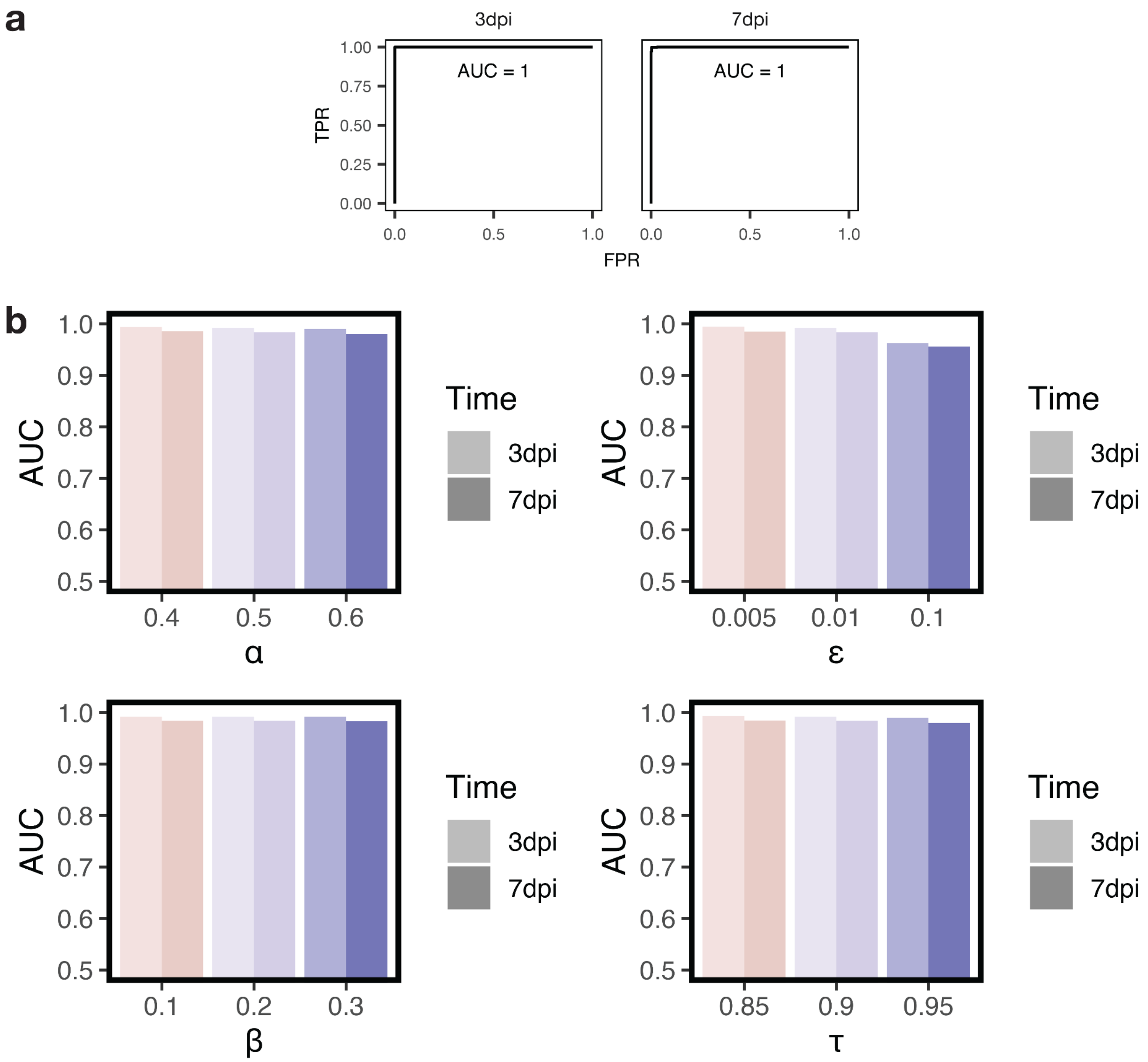


**Suppl. Fig. 11 | Moslin performance on zebrafish heart regeneration data is robust to variations in testing strategy and hyperparameter values.**

**a**. ROCs and AUCs for one-versus-rest cell type persistency test. **b**. Moslin performance on the zebrafish heart regeneration dataset as measured through the cell type persistency test is uniformly high at various hyperparameter values. Increasing $\epsilon$ to 0.1 reduces performance by 3% (from AUC 0.99 to 0.96 at both time points); variations in other hyperparameters yield performance changes below 1%. Baseline hyperparameters: $\alpha = 0.5$, $\beta = 0.2$, $\epsilon = 0.01$ and $\tau_a = 0.9$ (Methods).

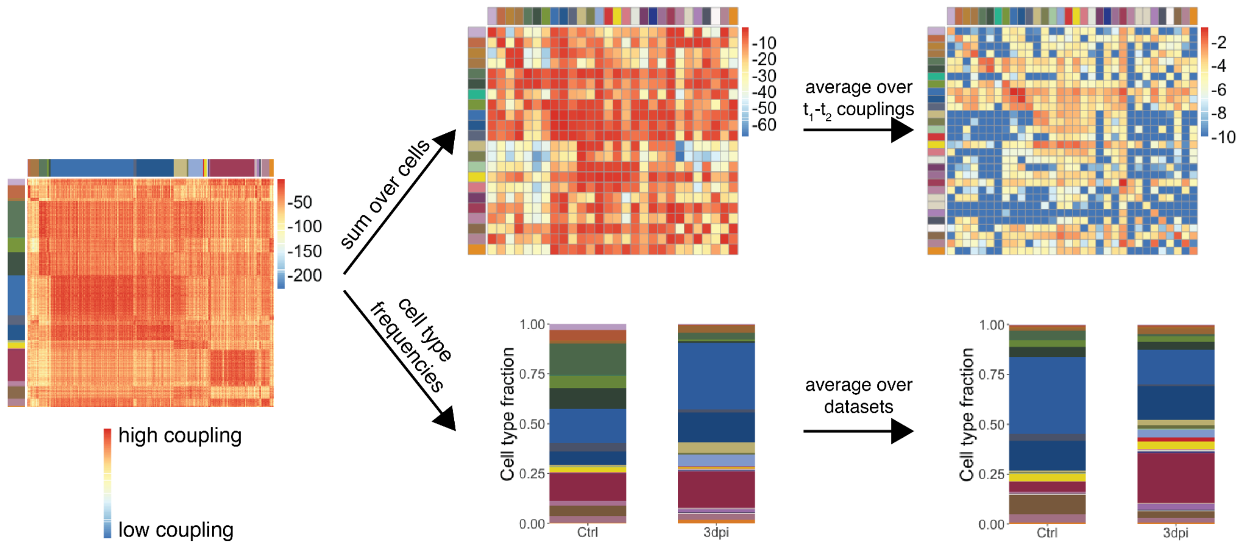


**Suppl. Fig. 12 | Calculation of cell type transitions in zebrafish heart regeneration.**

The moslin-calculated cell-cell coupling matrix for every combination of $t_1$ and $t_2$ datasets is aggregated to obtain cell type couplings; we also calculate cell type fractions. Both of these are averaged - the cell type fractions over all datasets and the couplings over all $t_1$-$t_2$ dataset combinations, and the averages are used to calculate cellular flows. Coupling color-scales shown are log-10 of the coupling values.
